# Molecular plasticity in the flavin binding pocket of BLUF domain evolves first light-gated endonuclease in bacterial system

**DOI:** 10.1101/2025.05.05.652191

**Authors:** Rahamtullah, Jitender, Manish Singh Kaushik, Sagar Kumar, Alfons Penzkofer, Andreas Möglich, Suneel Kateriya

## Abstract

The bacterium *Rubellimicrobium mesophilum* possesses a BLUF coupled endonuclease III (BLUF-EndoIII) with potential endonuclease activity. Interestingly, the crucial amino acid residues (tyrosine, histidine and tryptophan) responsible for BLUF photocycle and photodynamics are evolutionarily replaced by phenylalanine (Y5F), asparagine (H27N) and alanine (W87A) residues, respectively. In present communication, we have studied the impact of this evolutionary plasticity on the BLUF photodynamics and associated endonuclease activity. The results obtained showed that the evolutionary plasticity in the BLUF domain influenced various functional aspects of BLUF domain including FAD binding, domain stability, recovery kinetics, and spectral characteristics. The impact of amino acid plasticity on the C-terminal endonuclease (EndoIII) domain was also studied. The evolutionary plasticity induced changes in the flavin binding pocket of the BLUF domain elevated the light-gated endonuclease activity associated with EndoIII domain. The molecular docking analysis and spectroscopic studies also confirmed the substrate-binding ability of the BLUF-EndoIII. The elevated endonuclease activity suggested that the amino acid residues, which are crucial for BLUF photocycle are indeed dispensable and there might exists another electron transfer pathway for BLUF domain activation and regulation of associated endonuclease domain. Considering the role of endonucleases in bacterial defense, the understanding of the BLUF photodynamics, mechanism of signal transfer to the downstream endonuclease domain and associated endonuclease activity might elucidate the first naturally occurring light-gated endonuclease in bacterial system.

## Introduction

Microorganisms respond to the varying light conditions using variety of photoreceptors which execute light-driven control of associated output domains (Purcell and Crosson 2008). Blue-light using flavin (BLUF) proteins are the blue light responding photoreceptors, which upon illumination causes structural rearrangements around the chromophore to modulate the communion between BLUF and effector domains (enzymes or transcriptional regulators) to generate the full range of regulated photo-adaptive responses (Kaushik et al. 2019; Kennis and Mathes 2013; Barends et al. 2009). The mesophilic soil bacterium, *Rubellimicrobium mesophilum* strain MSL-20^T^ harbors a naturally occurring photoreceptor coupled restriction endonuclease, BLUF-Endonuclease III (hereafter BLUF-EndoIII) (Penzkofer et al. 2016). With a brief expansion of the C-terminus, the BLUF-EndoIII is made up of a BLUF domain and an effector domain which belongs to the endonuclease III family of DNA repair enzymes (Rao et al. 2014). In BLUF-EndoIII, the conserved flavin binding amino acids i.e. tyrosine, histidine and tryptophan (Y21, H44 and W104 in AppA protein) crucial for BLUF photocycle (Zhuo et al. 2024; Karadi et al. 2020; Anderson et al. 2005) have been naturally replaced by phenylalanine (F5), asparagine (N27) and alanine (A87), respectively (Penzkofer et al. 2016). In our previous study, we have performed the side-directed mutagenesis in BLUF-EndoIII and formed a triple mutant which was named as RmPAE (photo-activated endonuclease from *Rubellimicrobium mesophilum*) (Penzkofer et al. 2016). In RmPAE, the typical flavin binding pocket necessary for the BLUF domain photocycle has been restored. However, unlike other BLUF proteins, smaller spectral red-shift and slower signaling state recovery to the receptor state was observed in RmPAE (Penzkofer et al. 2016). The above observation suggested that that the evolutionary changes in the flavin binding pocket of BLUF domain of the BLUF-EndoIII might be enzymatically favorable. Therefore, to understand the implications of this evolutionary plasticity in the flavin binding pocket on the photocycle and photoactivation of BLUF domain, we have commercially synthesized a codon optimized sequence of RmPAE for the expression in *E. coli*. Thereafter, amino acids Tyr5, His27 and Trp87 were simultaneously mutated back to Phe5, Asn27 and Ala87, respectively; and the triple mutant (BLUF-EndoIII Y5F, H27N and W87A) obtained was compared with RmPAE (considered as wild type for this study) to examine the amino acid plasticity in relation to the photodynamic and photophysical features of the full-length BLUF domain and its role in controlling associated endonuclease activity.

The bacterial systems possess restriction-modification (R-M) system as an innate immune system to protect their genomes from the foreign invaders like bacteriophages (Lau et al. 2020; Nagamalleswari et al. 2017). The typical R-M system consists of endonucleases which act against all foreign and improperly-modified DNA. However, to match pace with fast-evolving anti-restriction strategies of bacteriophages, the bacterial defense also devised new mechanisms to counter phage infections (Labrie et al. 2010). The BLUF-EndoIII form *R. mesophilum* present one such mechanism where endonuclease domain is present in modular architecture with BLUF domain, hence; may be controlled by light. Several studies have already been performed in the past to precisely control the recombinant endonuclease activity by using external light signals (Li CY et al. 2022; Schierling et al. 2010; Zaremba and Siksnys 2010). However, BLUF-EndoIII provides researcher an interesting model which may be controlled endonuclease activity temporally without performing additional genetic engineering. The light-dependent endonuclease activity as well as the DNA binding ability of TM (representative of naturally occurring BLUF-EndoIII) were also evaluated in this study. Understanding the mechanism of BLUF photocycle and endonuclease domain activity in BLUF-EndoIII might elucidate the first naturally occurring light-gated endonuclease in bacterial system.

## Results & Discussion

### Homology analysis and biochemical characterization of RmPAE and TM

The identified BLUF-EndoIII from a pink to light reddish-pigmented mesophilic bacterium *R. mesophilum* strain MSL-20^T^, consists of a N-terminus BLUF domain (∼88 amino acids) and a C-terminus endonuclease domain (∼59 amino acids) (Fig. S1). Homology analysis of BLUF domain of BLUF-EndoIII from *R. mesophilum* with the well characterized BLUF domain containing proteins showed that most of the flavin binding residues are well conserved in BLUF-EndoIII (Fig. S2). However, conserved residues tyrosine (Y5), histidine (H27), and tryptophan (W87) (T21, H44, and W104 in AppA BLUF), were naturally replaced by phenylalanine (F), asparagine (N) and alanine (A), respectively; in the BLUF-EndoIII from *R. mesophilum* (Fig. S2). The highly conserved amino acids i.e. tyrosine (Y), asparagine (N), glutamine (Q), and tryptophan (W) or methionine (M) are crucial for the photodynamics and photocycle of the BLUF domains (Kaushik et al. 2019; Tanwar et al. 2016). The conserved tyrosine and glutamine residues in flavin binding site of the BLUF domain are used to sense light by mechanism involving rotamer shift of the glutamine side chain, which resulted into disruption in the hydrogen bonding network around the chromophore (Ohki et al. 2016). Upon blue light illumination, the rotated glutamine side chain interacts with Trp, and thereby causing structural changes in the BLUF protein (Fujisawa and Masuda, 2018). In dark state, the conserved glutamine involves in the hydrogen bond formation, whereas in light adapted conditions, the hydrogen bond is shifted to the conserved tyrosine via the process of a tautomerism (Kennis and Mathes, 2013).

Furthermore, multiple sequence alignment of endonuclease domains from naturally occurring BLUF-EndoIII was performed against well characterized MutY from *Escherichia coli* and other type III endonucleases (Fig. S3). The sequence alignment among the endonucleases reveals the presence of highly conserved 19 amino acid (aa) helix-hairpin-helix (HhH; Asp135-His157) motif in the endonuclease domain (Val113-Ala172) of BLUF-EndoIII, which is a characteristic feature among the DNA repair enzymes (Kanchan et al. 2015; Thayer et al. 1995; Fig. S3). The HhH motif within the endonuclease domain is responsible for binding protein non-specifically to the phosphate group of DNA via hydrogen bonding (Kanchan et al. 2015). The residues from Asp135-Ala140 and Arg146-Leu157 form the Helix1 (α1) and Helix2 (α2), respectively; in the HhH motif (Fig. S3). The 5 aa (Leu141-His145) interhelical domain forming β-turn which might act as the binding site for free thymine glycol, and hence the overall HhH motif may be considered as a DNA binding motif (Thayer et al. 1995). The hydrophobic amino acids Leu138, Leu141, Ile152 and Leu153 help to place the H1 against H2 and help stabilizing the HhH motif (Thayer et al. 1995). Further, the active site residue, aspartate (D165) (D138 in *E. coli* Endonuclease III) was also found conserved, which is catalytically essential for the endonuclease activity (Fig. S3). D165 helps in initiating the nucleophilic attack crucial for the glycosylase activity (Kanchan et al. 2015; Manuel et al. 2004). The presence of characteristic DNA binding HhH motif and conservation of active site residue suggests that the endonuclease domain in BLUF-EndoIII may belongs to the Endonuclease III family.

### Spectroscopic characterization of RmPAE and triple mutant

#### Blue light (BL) absorption photocycle and photodynamics of RmPAE and TM

The heterologous expression of recombinant proteins of RmPAE (wild type) and triple mutant (TM, BLUF-EndoIII Y5F, H27N and W87A) were induced in *E. coli*. The Recombinant proteins from RmPAE and TM were expressed as an N-terminal histidine (His) tag fusion proteins (5XHis:RmPAE and 5X His:TM), respectively. Recombinant proteins from RmPAE and TM were isolated using IMAC and resolved on SDS-PAGE). The SDS-PAGE profile of both RmPAE and TM showed a band of ∼28 kDa, which is known to be an estimated size of full-length BLUF Endonuclease IIIc (data not shown). The spectroscopic analysis suggested the functional expression of the purified recombinant proteins from RmPAE and TM in *E. coli* (Fig. 1A and B). The UV-visible absorption spectroscopy of purified recombinant proteins from RmPAE and TM was carried out between 300-700 nm wavelength range. The UV-visible spectra of purified recombinant proteins from RmPAE showed typical spectrum with vibronic peaks in the dark-adapted state with a maximum absorbance at 370 and 440 nm (Fig. 1A). Vibronic structures were observed at 420 and 470 nm, which is typical for flavin bound BLUF proteins (Fig. 1A). In case of wild type RmPAE, upon illumination with blue light, an ∼10nm red-shift in the spectrum was seen (Fig. 1A). Difference spectra between the dark-adapted and signaling state spectra showed maximum peak at 480 nm (Fig. 1A). The absorption spectrum for the TM was almost similar to that of the wild type RmPAE with absorption maxima at 374 and 440 nm (Fig. 1B). The typical vibronic shoulders at 416 and 460 nm were also well resolved (Fig. 1B). However, unlike RmPAE, a red-shift of ∼10 nm in the absorption spectra was not seen in TM (Fig. 1B). Upon blue light illumination of the BLUF domain, a photochemical intermediate with oxidized FAD is formed which is actually responsible for the ∼10 nm red-shift in the protein spectra (Iwata et al. 2018). An ultrafast transient spectroscopy of PixD demonstrated the involvement of Tyr, Gln, Trp and FAD in the formation of the red-shifted intermediate (Iwata et al. 2018). Therefore, mutation of the crucial amino acid Tyr (Y5) and Trp (W87) in TM might be responsible for the absence of the red-shift in the protein spectra.

**Fig. 1.**
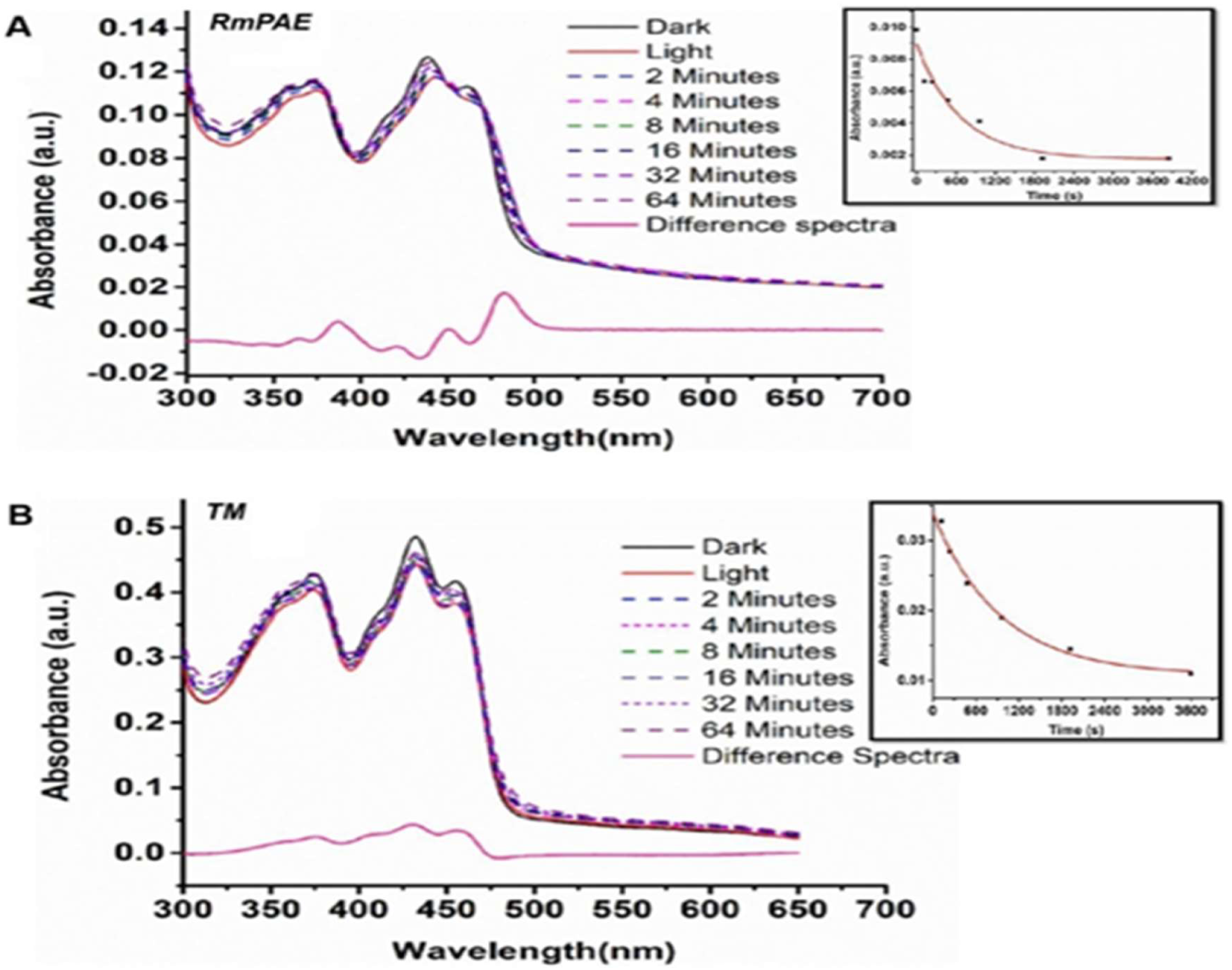
Spectral characterization of purified recombinant proteins from RmPAE and TM. (A) Absorption spectra of dark and light adapted RmPAE protein and the difference spectra. Dotted traces spectra acquired at different time intervals during the recovery of the protein in the dark. The time course of the decay of the UV-visible absorption changes at 440 nm induced by light in the RmPAE (inset image). (B) Absorption spectra of dark adapted and light adapted triple mutant BLUF endonuclease protein and the difference spectra. Dotted traces spectra acquired at different time intervals during the recovery of the protein in the dark. The time course of the decay of the UV-visible absorption changes at 440 nm induced by light in the triple mutant (TM; inset image).

Although, the spectral shift was not observed in TM, but it exhibited the photoexcitation following the blue light illumination (Fig. 1B). The dark state recovery following the illumination with blue light were observed for both RmPAE and TM. The signaling state of RmPAE reverts back to the dark ground state; however, the TM did not recover completely (Fig. 1A and B). The recovery rate analysis showed an increase in a time constant (τ = 687 s) for protein decay rate in TM when compared to that of RmPAE (τ = 462 s) in dark state (inset of Fig. 1A and B and Table S1). In BLUF proteins, blue light illumination causes photo-excitation which drives the transfer of electrons from Tyr residue to adjacent flavin. The electron transfer resulted into rearrangement of the H-bonds in the flavin binding pocket between N5 of the isoalloxazine ring and amino acid residues, which is responsible for the red absorption shift and formation of signaling state of the BLUF photoreceptor (Tanwar et al. 2016). In case of TM, the partial recovery in dark ground state was may be due to the possible changes occurred in the H-bonding environment in the flavin binding regions due to the loss of amino acids (Y5, H27 and W87) crucial for the typical BLUF photocycle. The photo-spectroscopic studies with SUMO-bPAC-Y7F also discussed the partial recovery of signaling state to the dark ground state (Penzkofer et al. 2014). Mutation of tyrosine to phenylalanine (Y5F), the key electron donor located in the vicinity of the flavin cofactor in the BLUF domain break the photo-induced electron transfer due to energetic reasons and changes the fate of flavin cofactor (Penzkofer et al. 2014; Mataga et al. 2002).

#### Fluorescence spectroscopic and decay lifetime study of RmPAE and TM

The fluorescent spectral analysis (kex=390 nm) of both RmPAE and TM in dark exhibited maximum emission peaks at 482 nm (Fig. 2A and B). Upon blue light illumination, a ∼10 nm spectral shift was observed both in RmPAE and TM (Fig. 2A and B). In BLUF proteins, blue light illumination causes fluorescence quenching due to increase in solvent exposure or disruption of π-π interaction, which is responsible for the ∼10 nm spectral shift (Zirak et al. 2006; Masuda et al. 2004; Kraft et al. 2003). The recovery kinetics of fluorescence spectra was almost coinciding with that of absorption spectra, where the signaling state recovery was observed in RmPAE protein, but not in the case of TM (Fig. 2A and B). The changes in fluorescence spectra following light illumination and its recovery in dark state suggests stability of the BLUF domain and functional photocycle. Therefore, loss of recovery kinetics in TM was may be disruption of the interactions in the flavin binding pocket of BLUF domain.

**Fig. 2.**
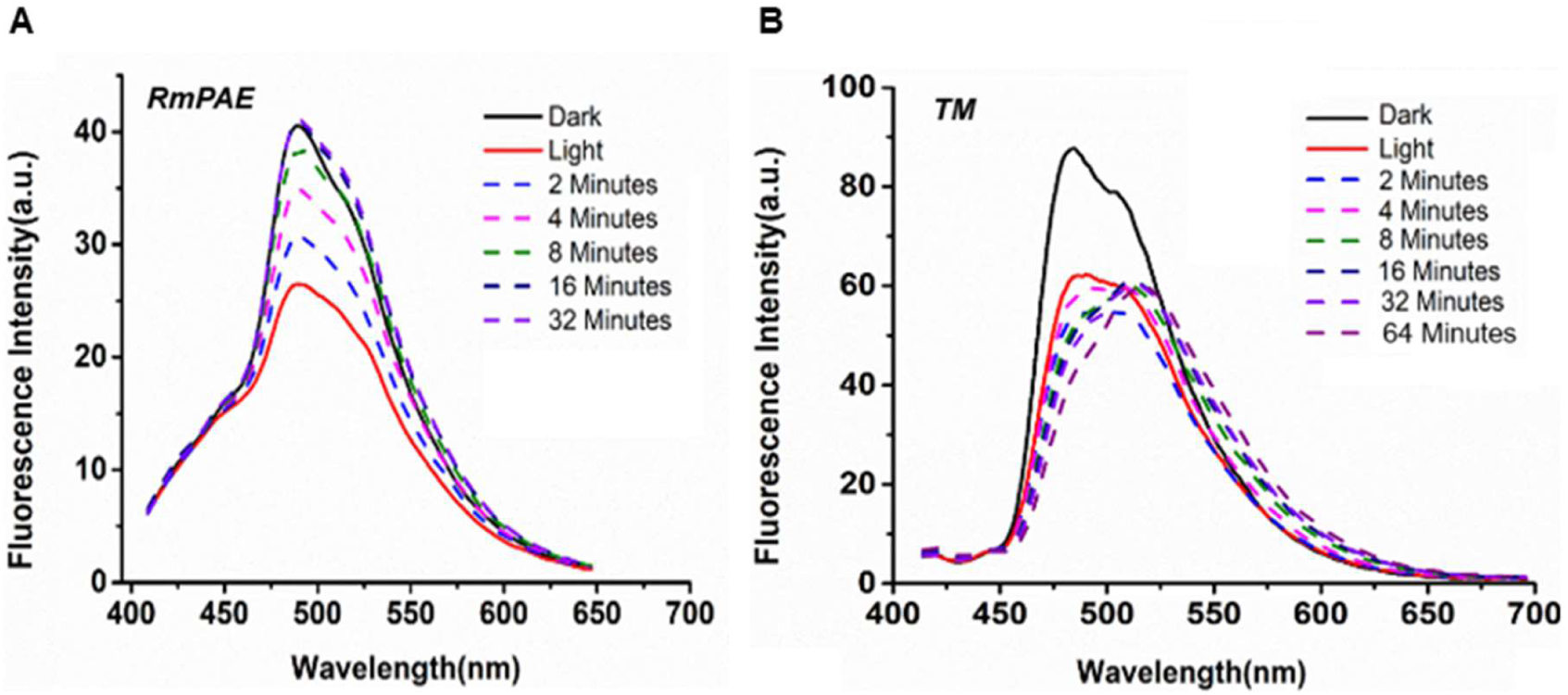
Fluorescence emission spectra of flavin from RmPAE (A) and TM (B), under dark and after exposure to blue light. Dotted traces are fluorescence emission spectra that were collected at various time points when the protein was recovering to the dark state. The sample were excited with 390 nm and scan range was 410-700 nm.

Furthermore, in order to ascertain the fluorescence lifespan of a fluorophore (FMN), time-resolved fluorescence spectroscopy (TRFS) was employed to examine the decay of the overall fluorescence intensity. The estimation of flavin’s fluorescence lifetime will provide important informations about the flavin surroundings. We have used the time-correlated single-photon counting (TCSPC) method for measuring fluorescence decay of RmPAE and TM (Fig. 3). For the RmPAE and TM proteins, three components were identified, each having a distinct lifetime and fractional values. The average fluorescence lifetime (<τ>) for the recombinant proteins from RmPAE and TM was determined to be 0.13 and 2.02 ns, respectively (Table S2). Flavoproteins upon light illumination exhibit ultrafast electron transfer (in picosecond and nanosecond time scales) from the aromatic amino acid residues (Trp and Tyr) to the electrically excited flavin chromophore which results in substantial shortening of the fluorescence lifetime of the flavin (Zhuo et al. 2024; Karadi et al. 2020). In BLUF proteins, tryptophan residue located in the proximity of flavin chromophore is responsible for the fluorescence quenching after blue illumination (Karadi et al. 2020). The Trp_in_NH_in_ conformation drives the one or two step proton transfer to flavin which resulted into fluorescence quenching by forming a Try-Gln configuration not conducive for proton transfer (Zhuo et al. 2024). In our case, mutation of tyrosine (Y5F) and tryptophan (W87A) residues in TM also resulted into an increment in the fluorescent lifetime, which suggests the loss of the typical electron transfer and tryptophan-dependent fluorescence quenching mechanism after photoexcitation with blue light. The exact mechanism for the enhanced fluorescence lifetime in the response of these mutations is still unknown and requires further studies.

**Fig. 3.**
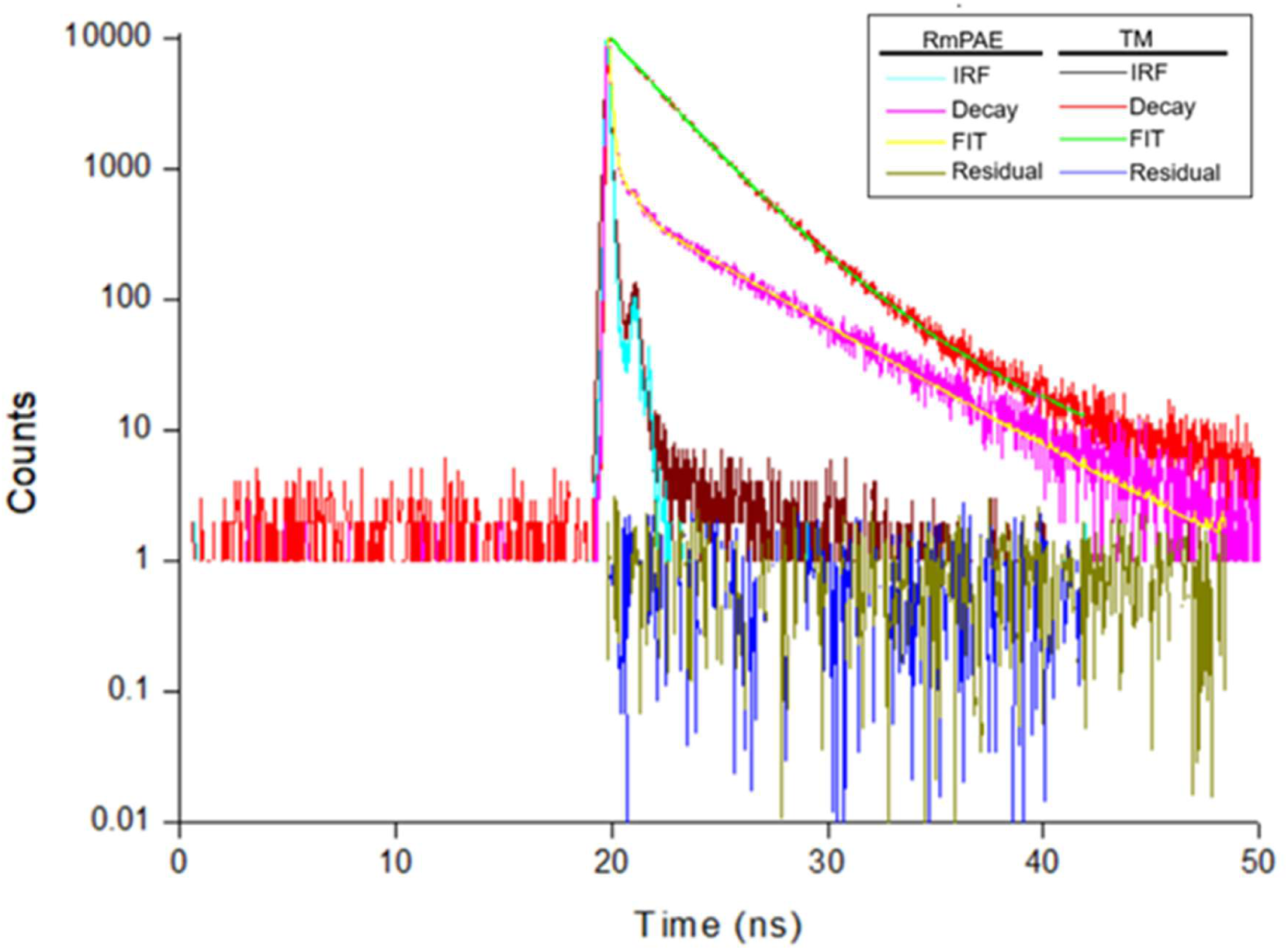
Time resolved fluorescence spectra (TRFS) for lifetime measurements of RmPAE (**A**) and triple mutant (TM; **B**). Data was fitted using three exponential functions.

#### Dynamics of light induced conformational changes in the RmPAE and TM

CD measurements were carried out in the far-UV (200–250 nm) range, in order to inquire the changes in the secondary structure of RmPAE and TM in dark and after blue light illumination. In both RmPAE and TM, a prominent negative ellipticity has been observed at 208 and 222 nm in dark state, indicates presence of α-helices, which is a characteristic feature of BLUF proteins (Fig. 4A and B). Dark CD spectra showed that both RmPAE and TM proteins were well folded and structurally similar (Fig. 4A and B). The RmPAE CD spectra showed a significant decrease in ellipticity after blue light illumination indicates light-induced conformational change in helical structures (Fig. 4A). The TM however exhibits almost negligible change in the ellipticity after blue light illumination, hence depict no conformational changes (Fig. 4B). The conserved amino acids tyrosine and tryptophan are involved in the strengthening of the hydrogen bond interaction with the flavin chromophore in the light-induced signaling state (Masuda et al. 2005). The spectroscopic studies with AppA showed that mutating tryptophan to alanine (W104A) causes conformational distortions by disturbing the hydrogen bonding interaction between Trp104, O4 of FAD and Gln63, crucial for the formation of the light-induced signaling state. Loss of tyrosin and tryptophan blocks the transformation of light signal into a specific β-sheet conformation in BLUF domain for the attainment of signaling state (Masuda et al. 2005). On the other hand, the histidine residue H27 (H44 in AppA) is specifically involved in transducing the light signal to the exposed surface of BLUF domain via active site residue Asn45 (Jung et al. 2006). Therefore, replacing crucial amino acids (Y5, H27 and W87) influence the hydrogen bonding interaction in the flavin bonding pocket, and hence no conformation of the secondary structure was observed in TM.

**Fig. 4.**
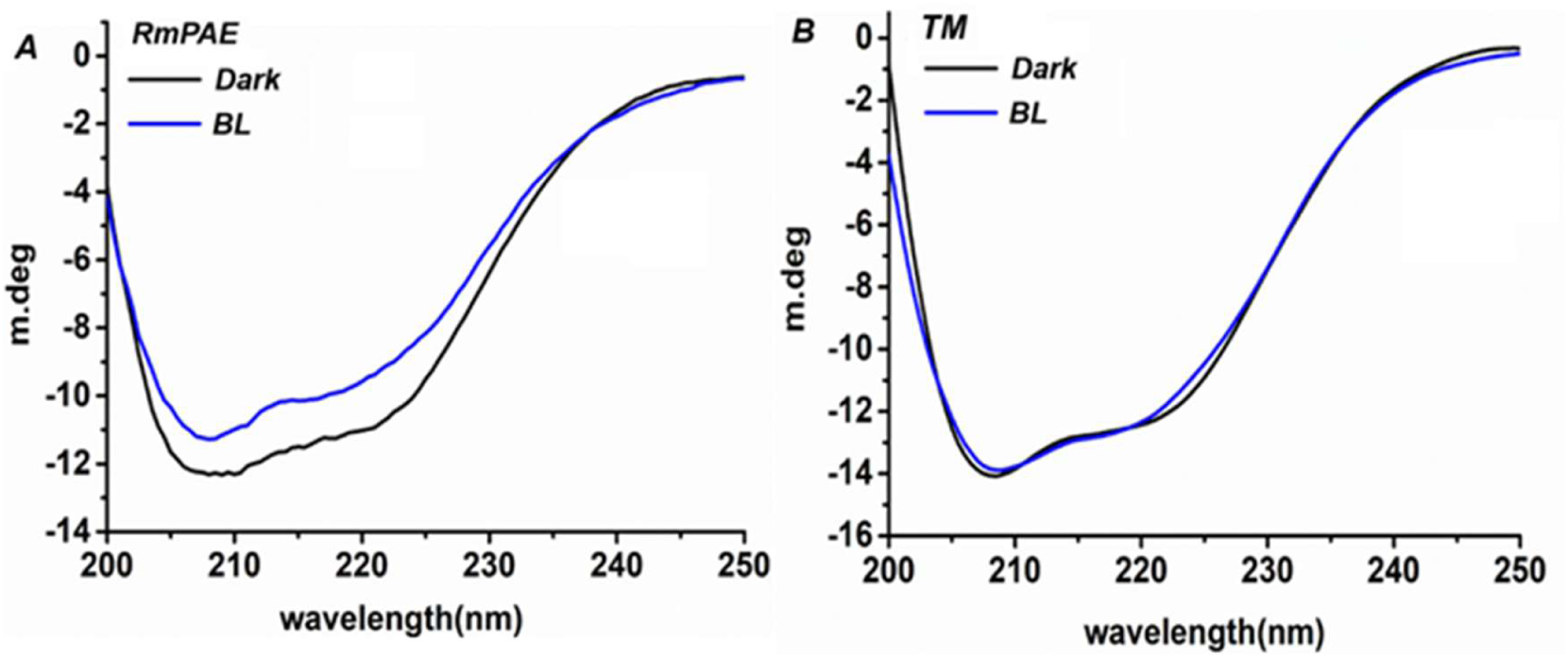
Secondary structure analysis of RmPAE and triple mutant (TM). Comparative CD spectra of the dark adapted and 30 second blue light (BL)-illuminated RmPAE (**A**) and triple mutant (TM) (**B**). The dark-adapted and BL-illuminated state of protein are represented as black and blue colour, respectively.

### 3D modeling of RmPAE and triple mutant and docking analysis with FAD

Tertiary structure prediction was carried out separately for the RmPAE and TM (Fig. S4A and B). The predicted 3D model of both RmPAE and TM showed the presence of a typical BLUF domain with five-stranded β-sheet flanked by two α-helices (Fig. S4A and B). The 3D model of RmPAE and TM also posses a helix-hairpin-helix (HhH) motif (Fig. S4A and B), which is a characteristic feature of members of endonuclease family (Thayer et al. 1995). The helix-hairpin-helix (HhH) motif consists of two α-helices i.e. α-helix 1 (α1) and α-helix 2 (α2), separated by a β-turn (Fig. S4A and B), which is in the coherence with our sequence analysis data (Fig.S3A). As we discussed earlier, like other well characterized endonuclease III, the β-turn in the HhH motif might act as the binding site for free thymine glycol (Thayer et al. 1995). Further, the docking analysis revealed that the FAD was readily accommodated in the flavin binding pocket sandwiched between two α-helices in the modelled BLUF domains; however, the protein-ligand interaction pattern was significantly changed when compared between RmPAE and TM (Fig. 5A-F). In RmPAE, Gln46 and Asn28 located near the isoalloxazine ring of FMN interact via forming hydrogen bond (Fig. 5A-C). In BLUF proteins, the FAD chromophore is non-covalently linked to Try, Gln and Asn residues via hydrogen bonding network (Gil et al. 2017). Upon photoexcitation, Gln interacts with a conserved tyrosin’s (Tyr8) phenolic hydroxyl via forming hydrogen bond, resulting in a FAD-Gln-Tyr network (Gil et al. 2017). Similarly, in AppA protein, FAD chromophore forms H-bonds with conserved residues Tyr21, Gln63 and Trp104 (Masuda et al. 2013). The observed red shifts in the spectra were proposed to be caused by the rotation of the glutamine side chain during photoexcitation. Due to this rotation and formation of new hydrogen bond with the flavin carbonyl group, the electronic cloud of the isoalloxazine ring of flavin gets altered, thereby giving rise to a ∼10 nm red shift in the absorption spectrum (Dragnea et al. 2005; Unno et al. 2006). The interaction between Gln and carbonyl group of FMN, which is crucial for the BLUF photocycle was found distorted in case of TM (Fig. D-F). The replacement of tyrosine (Y5F) and tryptophan (W87A) in TM might cause some structural changes, which resulted into placing Gln out of the hydrogen forming distance of the flavin carbonyl group. As we discussed earlier, the predicted tertiary structures of RmPAE and TM showed similar conformation of BLUF domain as well as the similar flavin binding pockets (except replaced amino acids i.e. Y5, H27 and W87) (Fig. S4A and B). However, when we overlayed the FAD-docked 3D models of RmPAE and TM, TM depicted conformational flipping of the BLUF-domain (Fig. 6). This interesting observation suggested that the replacement of above-mentioned amino acids not only distorted the interaction pattern in the BLUF domain, but also changes the orientation of the BLUF domain upon flavin binding. However, the exact mechanism involved in this conformational flipping is still unknown and requires further investigation.

**Fig. 5.**
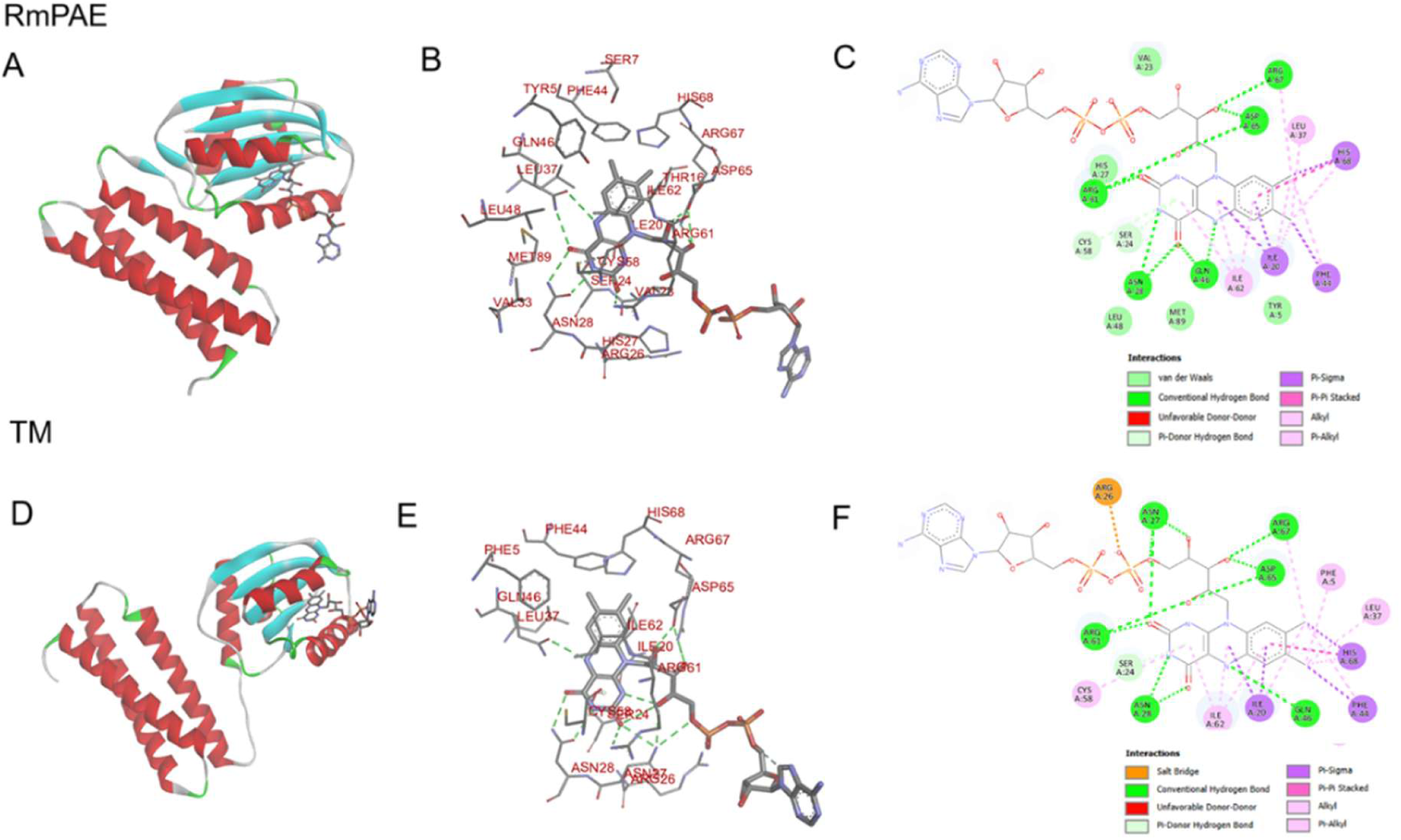
Molecular docking of FAD with BLUF domain of RmPAE (A) and TM (D). The FAD interacting amino acid residues and the type of interactions involved in the FAD binding in RmPAE (B, C) and TM (E, F).

**Fig. 6.**
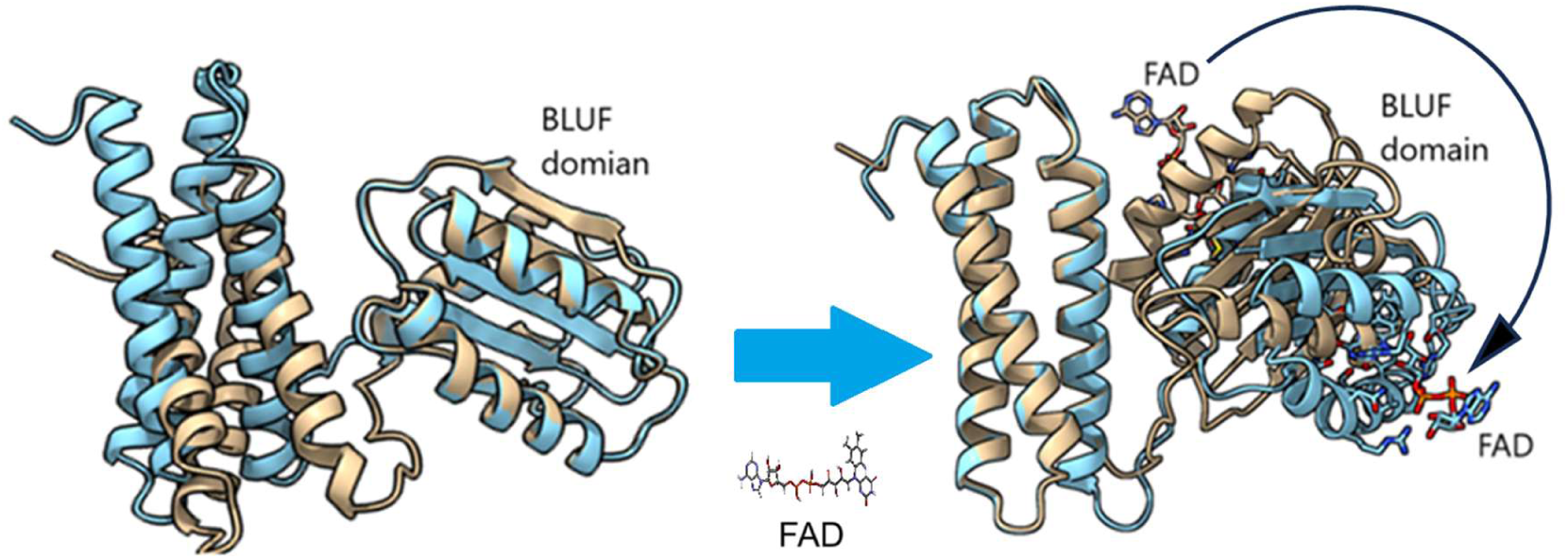
Superimposition of the tertiary structures of RmPAE (brown) and TM (cyan) in apo-(A) and FAD-bound (B) state. The FAD binding in the flavin binding site resulted into the flipping of the BLUF domain.

### Analysis of confirmational stability of RmPAE and TM

MD simulations were performed to study the structural changes caused by the mutations over the 100 ns run and the root mean square deviations (RMSDs) for the RmPAE and TM (with and without bound FMN) were assessed (Fig. S5A). The average RMSDs for RmPAE-apo, RmPAE in FMN-bound form, TM-apo, and TM in FMN-bound form were estimated 0.016 nm, 0.124 nm, 0.039 nm, and 0.152 nm, respectively. The RMSD plot showed that the RmPAE-apo form is comparatively more stable than the TM-apo form, as the RMSD plot of the later showed higher fluctuation in RMSD values near 30 ns (Fig S5A). Similarly, the RMSD plot of the FMN-bound TM was also less stable in comparison to the FMN-bound RmPAE as it shows an increasing RMSD value over the 100 ns simulation (Fig S5A). Therefore, above observations suggested that the recombinant protein from TM was comparatively unstable than that of recombinant protein from RmPAE, both in apo as well as in FMN-bound state. Further, the root mean square fluctuations (RMSFs) were also monitored during the MD simulation study to assess the residual flexibility in the BLUF domain of RmPAE and TM. The RMSF results indicated that the FMN-bound form of both RmPAE and TM behaved almost similarly (Fig. S5B); hence, suggesting the rigidity of the FMN-binding pocket. The radius of gyration (Rg) was also measured for the RmPAE and TM, both in apo and FMN-bound form (Fig. S5C). The Rg plot showed almost similar behavior both in RmPAE-apo and TM-apo. However, when the Rg plot for RmPAE and TM were compared in FMN-bound state, then the Rg values for the FMN-bound RmPAE was found lower in comparison to that of FMN-bound TM (Fig. S5C), suggesting that the compactness of the TM decreases in complex with FMN. The stability and compactness of the TM decreased due to loss of amino acid residues involved in interaction with the flavin.

We used principal components analysis (PCA) to gain a better understanding of the large-scale combined dynamics of BLUF domain conformation in RmPAE and TM, both in apo and FMN-bound form. PCA determines the dominant protein conformational patterns in a principal components (PCs) phase space during the 100 ns MD simulations. We projected the Cα atoms of the proteins along the direction of the first three eigenvectors (PC1 and PC2) to study their conformational behavior in the BLUF domain of RmPAE and TM in the presence of FMN. The initial three PCs for a given protein, taken from their corresponding 100 ns MD simulation trajectories, formed cluster groups. A 2D principal component plot was created to examine allowable conjoined motions between eigenvectors 1 and 2. Each dot in this 2D map represents a different trajectory conformation at a given time. The plot shows fluctuations in the ensemble distribution for each conformation throughout the simulation interval (Fig. S5D). The PCA plot shows that the collective motion of the RmPAE complex is relatively confined in a small conformation space, whereas the collective motion of the TM type occupies a broader conformational space, suggesting decreased complex stability in the TM. Additionally, the hydrogen bonding between BLUF domains of RmPAE and TM and the FMN were assessed, and the time progress of this bonding across the 100 ns simulation run is presented (Fig. S5E). The maximum number of hydrogen bonds in the TM was 5, whereas in the RmPAE was 4 (Fig. S5E). The hydrogen bonding (HB) plot shows that the TM forms slightly more hydrogen bonds than the RmPAE initially, but after 40 ns, it shows a similar pattern (Fig. S4E). We have also examined the Solvent Accessible Surface Area (SASA) of BLUF domains of RmPAE and TM to determine how the mutations impact the native protein’s structure. The SASA plot displays the SASA values determined over time for the BLUF domains of RmPAE and TM (Fig. S5F). The observations shows that the solvent accessible surface of BLUF domain in FMN-bound TM decreases as compared to the FMN-bound RmPAE, suggesting decreased stability of the mutant type. Overall, the outcomes of MD simulation suggested that the BLUF domain in TM has reduced stability in FMN-bound state.

### Photo-induced endonuclease activity in RmPAE and TM

We have assessed the endonuclease activity of RmPAE and TM by looking for DNA cleavage products visible on the agarose gel. The purified recombinant proteins of RmPAE and TM were incubated for 30 mins at 37 °C with substrate plasmid DNA (pGEX vector) in the absence (dark) and presence of blue light. After 30 mins of blue light illumination, the band representing the supercoiled form of the substrate plasmid was found to be absent both in RmPAE and TM, which suggests that the BLUF-coupled endonuclease domain has enzymatically cleaved the supercoiled form of the plasmid (Fig. 7A and B). However, in contrast to the RmPAE, where the intensity of the band representing the nicked or open circular form of plasmid has been increased, TM showed no reminiscence of plasmid DNA on the agarose gel after 30 mins of blue light illumination (Fig. 7A and B). This observation suggests two things, first the BLUF-coupled endonuclease initially digested the supercoiled form of plasmid followed by the nicked or open circular form and; secondly, the TM showed enhanced catalytic activity in comparison to that of RmPAE. We can assume that the BLUF domain regulates the nicking efficiency of endonuclease domain in a light-dependent manner. Further, when we compared the RmPAE and TM under dark condition, TM showed some catalytic activity in contrast to RmPAE where no endonuclease activity was observed (Fig. 7A and B). The observation showed that TM also showed some basal catalytic activity under dark condition which was enhance efficiently after blue light illumination under the influence of BLUF photocycle. The above observations suggest that the architecture of the flavin binding pocket of TM exhibited efficient DNA nicking ability in comparison to RmPAE.

**Fig. 7.**
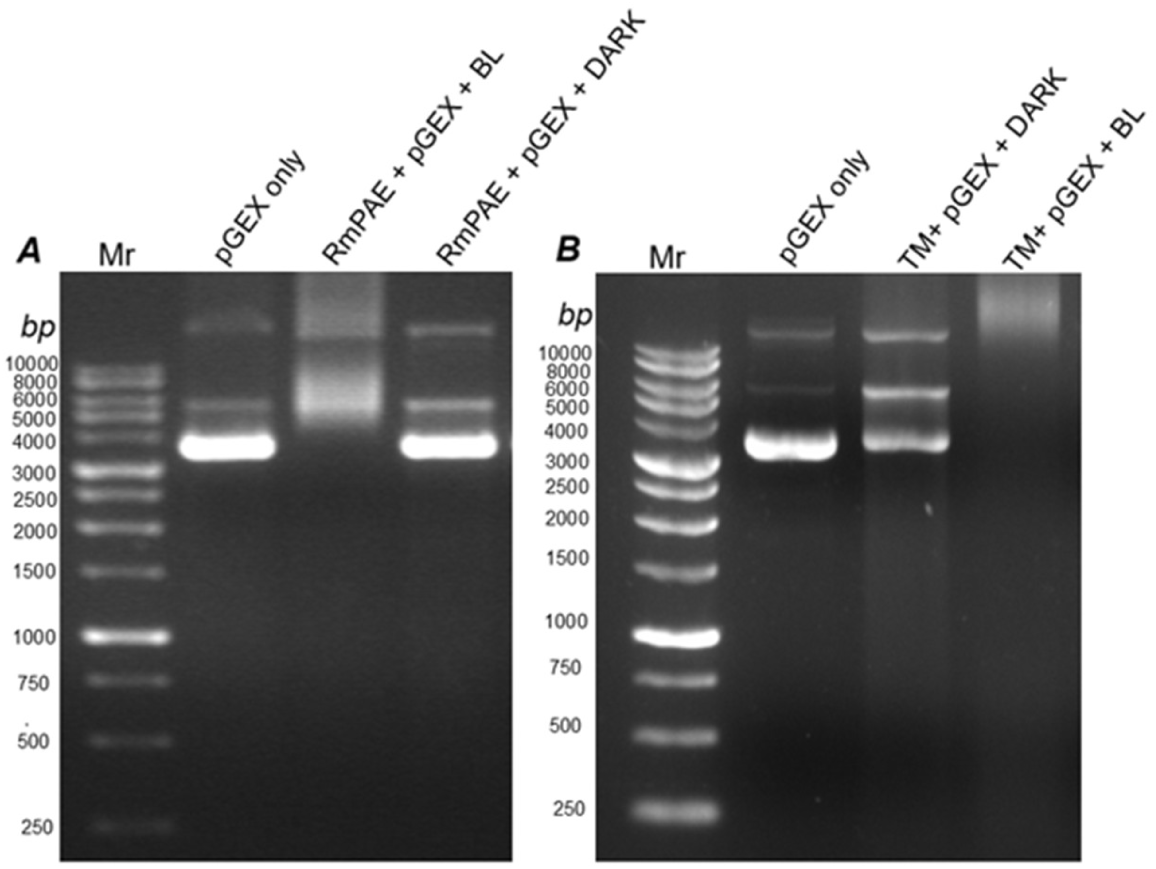
Endonuclease assay of purified recombinant RmPAE (**A**) triple mutant (TM; **B**). The constructs (pGEX; 500 ng) was incubated with purified RmPAE and TM protein (10µg) for 30 minutes in dark and after blue light irradiation at 37°C. Construct only was taken as the negative control.

### Triple mutant bind non-specifically to the DNA substrate

We performed the docking analysis by using TM and different fragments of plasmid DNA (pGEX vector; 500 bps each) as receptor and ligand molecule, respectively. The docking analysis revealed that TM bound non-specifically to three different plasmid DNA fragments (pGEX 1001-1500, pGEX 3001-3500 and pGEX 4501-4934) (Fig. 8A-C; Table S3). The TM-binding region on the plasmid DNA varies in length between 11-14 bp (Fig. 8A-C). The interaction between each plasmid DNA fragments and the TM was observed in the predicted EndoIII domain close to the HhH motif (Fig. 8A-C), which is considered as the DNA binding motif in the type III endonucleases family (Thayer et al. 1995). Asp 165, a conserved active site residue crucial for the endonuclease activity was found present among the amino acids interacting with the DNA fragment in each case (Fig. 8A-C; Table S3). Asp165 helps in initiating the nucleophilic attack crucial for the glycosylase activity (Kanchan et al. 2015; Manuel et al. 2004). To confirm the interactions, we have commercially synthesised each of the predicted binding sites and employed them for the DNA-protein interaction studies against the TM by monitoring the changes in the internal fluorescence. In comparison to unbound TM, a significant decrease in the fluorescence intensity was observed in oligonucleotide-bound TM at increasing concentration of oligonucleotide (Fig. 8D-F). The fluorescence quenching effect was monitored by estimating Stern-Volmer constant (K_sv_) and bimolecular quenching rate constant (K_q_) for each reaction (Fig. 8G-I; Table 1). The K_q_ values for all the three oligonucleotide-TM quenching reactions were higher than the maximum scatter collision quenching constant recorded for different quenchers-biopolymer complex i.e. 2.0×10^10^ M^−1^s^−1^ (Peng et al. 2015), which suggested the oligonucleotide dependent static quenching of TM fluorescence (Peng et al. 2015; Tian et al. 2012). The quenching of the fluorescence intensity confirmed the molecular association between the oligonucleotide and the TM. If we combined the outcomes of endonuclease assay and DNA protein interaction analysis, we could infer that TM may bound to the DNA substrate and cleave it by associated endonuclease activity.

**Fig. 8.**
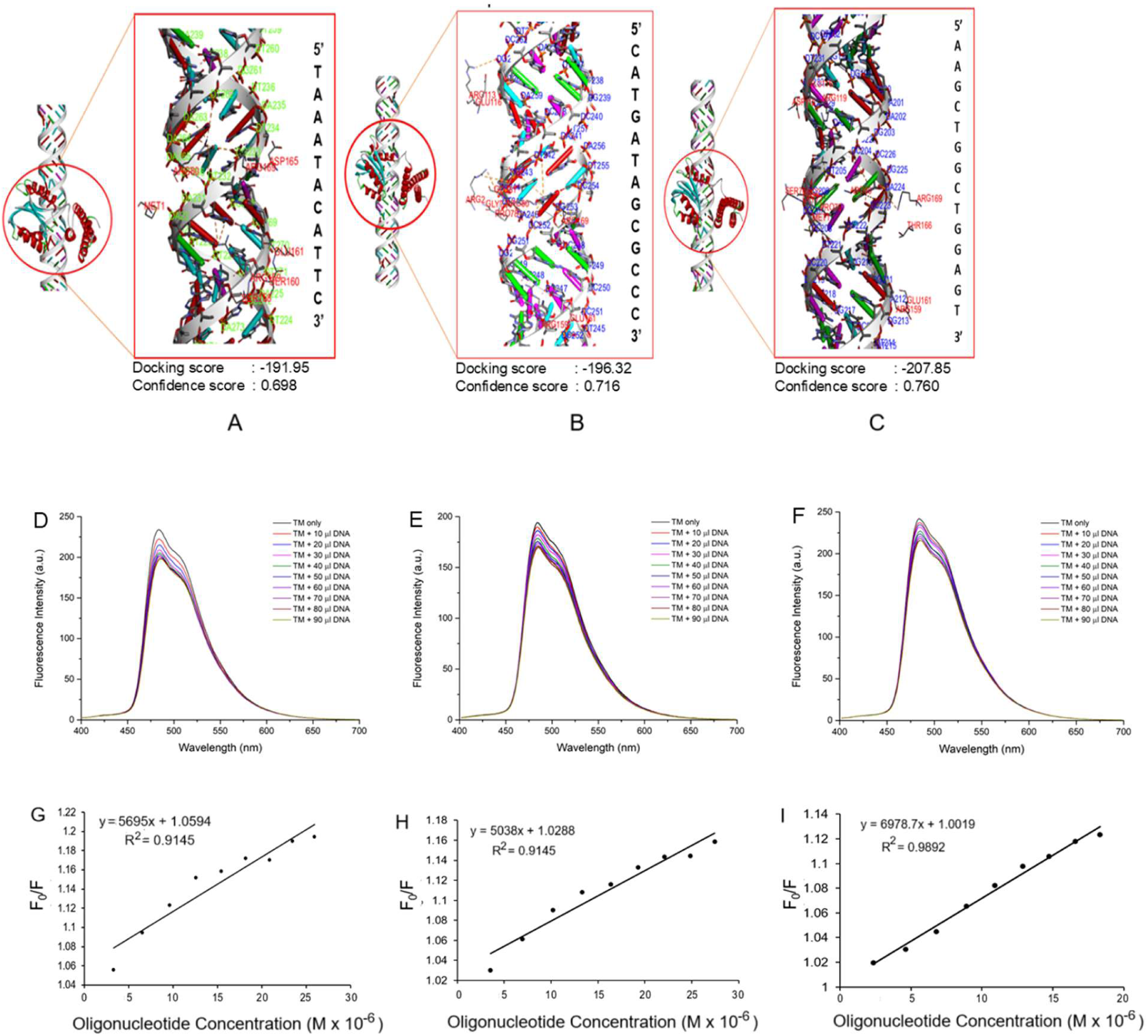
Recombinant TM and pGEX DNA interactions analysis. The docking analysis were performed by HDOCK server (http://hdock.phys.hust.edu.cn/) using TM and different fragments of pGEX vector [1001-1500 bps (A), 3001-3500 bps (B), and 4501-4934 bp (C)] as receptor and ligand molecules, respectively. The nonspecific binding sites of TM for each fragment is presented as 5’-3’ nucleotide segments. The length of nonspecific binding sites varies between 11-14 bps. DNA-dependent fluorescence quenching of recombinant TM (D-F). The Stern−Volmer plot depicting the oligonucleotide dependent fluorescence quenching of TM (G-I). The binding sites sequences predicted (Table S3) in fragments of pGEX vector [1001-1500 bps (D, G), 3001-3500 bps (E, H), and 4501-4934 bp (F, I)] were used for the oligonucleotide-TM interaction analysis. F_0_ – fluorescence intensity of unbound TM; F – fluorescence intensity of oligonucleotide-bound TM

**Table 1.** Stern-Volmer quenching constants (K_SV_) and bimolecular quenching rate constants (kq) of Oligonucleotide and protein complex (TM )at 37° C.

| Complex | $K_{sv}$ (M<sup>-1</sup>) | $K_q$ (M<sup>-1</sup>S<sup>-1</sup>) | $R^2$ |
| --- | --- | --- | --- |
| TM-pGEX vector 1001-1500 | 5.69 x 10<sup>3</sup> | 2.7 x 10<sup>12</sup> | 0.9145 |
| TM-pGEX vector 3001-3500 | 5.03 x 10<sup>3</sup> | 2.4 x 10<sup>12</sup> | 0.9462 |
| TM-pGEX vector 4501-4934 | 6.97 x 10<sup>3</sup> | 3.4 x 10<sup>12</sup> | 0.9892 |
| TM-RSPI01 1-500 | 15.4 x 10<sup>3</sup> | 7.6 x 10<sup>12</sup> | 0.9860 |
| TM-RSPI01 1501-2000 | 11.2 x 10 | 11.2 x 10<sup>12</sup> | 0.8984 |
| TM-RSPI01 2501-3000 | 9.5 x 10<sup>3</sup> | 9.5 x 10<sup>12</sup> | 0.9574 |

*TM* – triple mutant; *RSPI01*- Roseophage *SI01*; $K_{sv}$ – stern-volmer quenching constant; $K_q$ – bimolecular quenching rate constant; $R^2$ – correlation coefficient

To check the potential of BLUF-EndoIII as an optically-switchable bacterial immunity system, we have also performed the docking analysis between TM and DNA fragments of *Roseophage* SI01 (RSPI01) as receptor and ligand molecule, respectively. We select RSPI01, a lytic marine phage which infect bacteria belong to Rodobacteraceae family, which also include *R. mesophilum.* Like pGEX fragments, TM also bound non-specifically to different RSPI01 DNA fragments (RSPI01 1-500, RSPI01 1501-2000 and RSPI01 2501-3000) (Fig. 9A-C; Table S4). In case of RSPI01, TM-binding region varies in length between 11-13 bp (Fig. 9A-C). Like in case of pGEX fragments, the RSPI01 oligonucleotide-TM interactions were also observed in the predicted EndoIII domain close to the HhH motif with Asp 165 present among the interacting amino acids in each case (Fig. 9A-C; Table S4). Further, we have commercially synthesised sequences of each of the predicted binding sites in the RSPI01 oligonucleotide fragments and employed them for the DNA-protein interaction studies using fluorescence spectroscopy. The changes in internal fluorescence were monitored as a confirmation of oligonucleotide-TM binding (Fig. 9D-F). A significant decrease in the fluorescence intensity was observed in oligonucleotide-bound TM at increasing concentration of RSPI01 oligonucleotide fragments in comparison to the unbound TM (Fig. 9D-F). The estimated K_sv_ and K_q_ values for each reaction are given in Fig. 9G-I and Table 1. Like pGEX-TM interaction, the K_q_ values for all the three oligonucleotide-TM quenching reactions were also higher than 2.0 × 10^10^ M^−1^s^−1^, which suggested the RSPI01 oligonucleotide dependent static quenching of the TM fluorescence (Peng et al. 2015; Tian et al. 2012). The oligonucleotide-dependent quenching of TM fluorescence confirmed the bimolecular association between TM and phage oligonucleotide fragments. The binding of the RSPI01 oligonucleotide fragments in the predicted DNA binding motif of EndoIII domain close to the HhH motif suggested that these fragments might act as substrate for the TM.

**Fig. 9.**
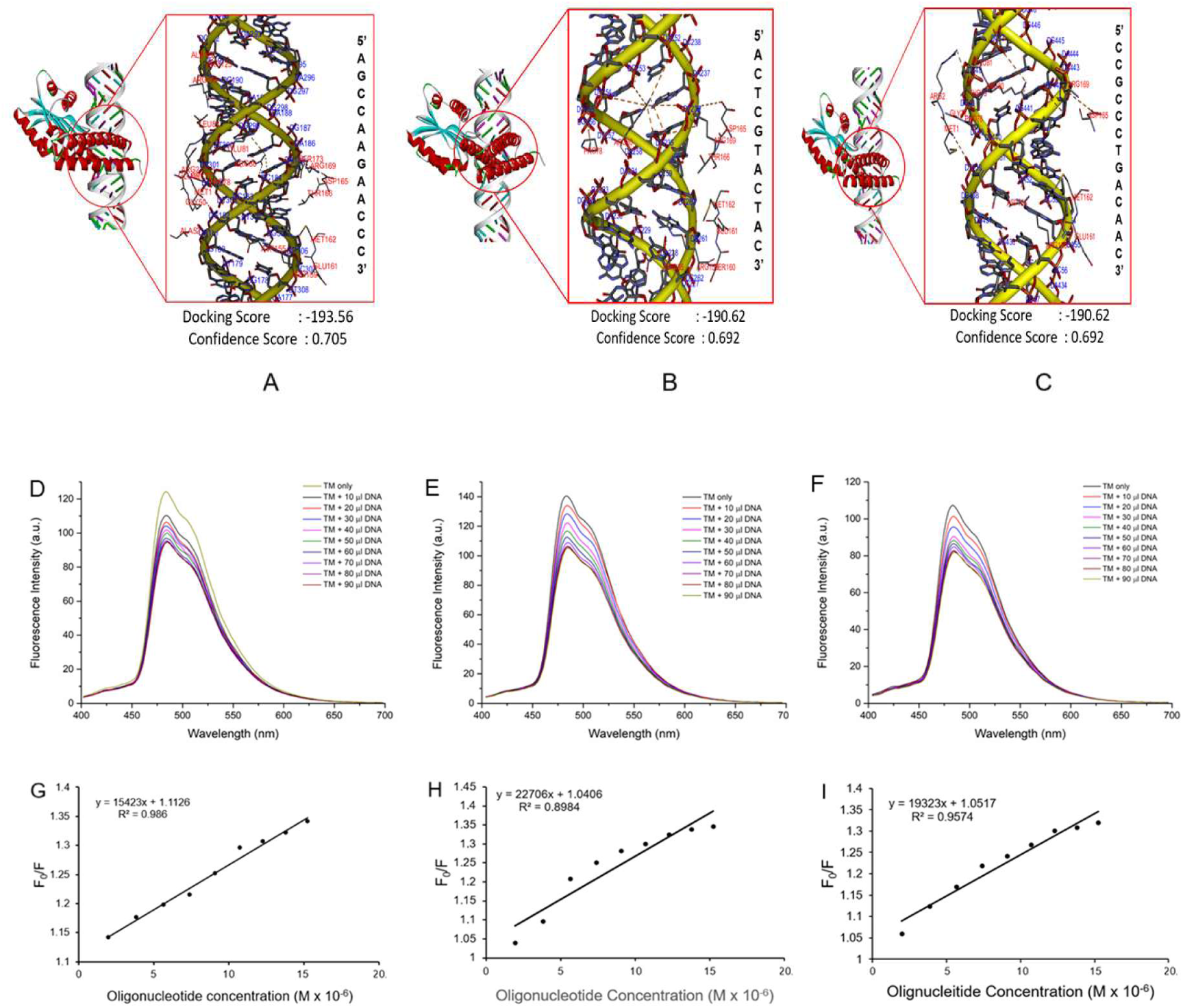
Recombinant TM and phage DNA interactions analysis. The docking analysis were performed by HDOCK server (http://hdock.phys.hust.edu.cn/) using TM and different fragments of *Roseophage* SI01 (RSPI01) DNA [1-500 bps (A), 1501-2000 bps (B), and 2501-3000 bp (C)] as receptor and ligand molecule, respectively. The nonspecific binding sites of TM for each fragment is presented as 5’-3’ nucleotide segments. The length of nonspecific binding sites varies between 11-13 bps. DNA-dependent fluorescence quenching of recombinant TM (D-F). The Stern−Volmer plot depicting the oligonucleotide dependent fluorescence quenching of TM (G-I). The binding sites sequences predicted (Table S4) in fragments of RSPI01 [1001-1500 bps (D, G), 3001-3500 bps (E, H), and 4501-4934 bp (F, I)] were used for the oligonucleotide-TM interaction analysis. F_0_ – fluorescence intensity of unbound TM; F – fluorescence intensity of oligonucleotide-bound TM.

## Conclusions

In this study, we have assessed the impact of evolutionary plasticity in the flavin binding pocket of BLUF domain on the BLUF photocycle as well as on the associated endonuclease activity. We have performed the comparative spectral analysis of RmPAE and TM in dark and after blue light illumination. Absorption spectra in dark showed similar spectral properties in RmPAE and TM. Upon blue light illumination, both RmPAE and TM exhibited ∼10 nm red shift in the spectra; however, unlike RmPAE, the spectral recovery was not observed in TM. The fluorescence emission spectra and the fluorescence recovery kinetics of RmPAE and TM showed similar behavior as observed in absorption spectra analysis, where the loss of signaling state recovery was observed in the case of TM. The TM also showed the decrement in the fluorescence decay and fluorescence lifetime in comparison to the RmPAE. The CD spectra analysis further showed no conformational change in TM upon blue light illumination in contrast to the RmPAE where significant conformational change was revealed. The loss of spectral recovery to the dark ground state, absence of conformational changes and decrease in fluorescence lifetime in TM was may be due to the mutation of amino acid residues (Y5F, H27N and W87A) crucial for the BLUF photodynamics. The MD simulation analysis also showed that the replacement of crucial amino acid residues in TM significantly influenced the confirmational stability, residual flexibility, compactness of flavin binding pocket and hydrogen bond interactions. The BLUF-coupled endonuclease domain from both RmPAE and TM exhibited light-dependent plasmid nicking ability; however, enhanced catalytic activity of endonuclease was observed in TM. The BLUF-coupled endonuclease preferentially digested the supercoiled form of plasmid followed by the open circular form. Interestingly, unlike RmPAE, TM also showed basal level catalytic activity in dark which was enhanced efficiently after blue light illumination under the influence of BLUF photocycle. The above observations suggested that the evolutionary plasticity in the flavin binding pocket of BLUF domain in TM although distorted the typical BLUF photodynamics, it might be activating an alternate pathway which promotes efficient DNA nicking ability in comparison to RmPAE. However, further analysis is anticipated to support this hypothesis.

The bioinformatics and spectroscopic analysis confirm the interaction between TM and oligonucleotide fragments. TM bind nonspecifically to the DNA binding sites of varying lengths. The interaction between TM and phage (RSPI01) oligonucleotide fragments were also showed by performing bioinformatics and spectroscopic analysis. The TM-phage DNA fragment interaction extrapolate the potential of this photoactivated endonuclease in bacterial defense against invading phages.

## Materials & Methods

### Domain identification and homology analysis of BLUF-EndoIII

The domain architecture of BLUF-EndoIII from mesophilic bacterium *R. mesophilum* strain MSL-20^T^ was analyzed using conserved domain retrieval tool (CDART; http://www.ncbi.nlm.nih.gov/Structure/lexington/lexington.cgi) (Geer et.al 2002). Homology analysis of the sequences of BLUF domain from RmPAE (codon optimized, commercially synthesized and wild type for this study), triple mutant (TM) and naturally occurring BLUF-EndoIII from *R. mesophilum* was carried out with the well characterized BLUF domain of AppA protein from *Rhodobacter sphaeroides*. Similarly, the homology analysis of endonuclease domains from RmPAE and naturally occurring BLUF-EndoIII was performed against well characterized MutY from *Escherichia coli* and endonuclease III domains from *E. coli, Micrococcus luteus*, and *Methanothermobacter thermautotrophicus,* respectively. The sequence-based homology analysis of BLUF and endonuclease domains was performed using BioEdit v7.2 (Hall, 1999)

### Site-directed mutagenesis, heterologous expression and purification of recombinant RmPAE and triple mutant

The Codon optimized sequence of RmPAE with typical amino acids (Y5, H27 and W87) crucial for BLUF photodynamics was synthesized for expression in *Escherichia coli* from BIOLINKK, New Delhi, India. The synthesized gene was cloned in pASKIBA43(+) vector (Novagen Inc, Madison, WI, USA) and point mutations (Y5F, H27N and W87A) were carried out using Quick change mutagenesis kit (Agilent Technologies, Santa Clara, CA, USA) as per manufacturer’s instructions. All constructs were sequenced, and mutations were confirmed. Proteins were expressed in *E. coli* BL21 (λDE3) and the recombinant proteins were purified using immobilized metal affinity chromatography (IMAC). Further, the purified protein samples prepared in Laemmle buffer were separated on 12 % SDS-PAGE. Then purified proteins were used for further spectroscopic characterization and endonuclease assay.

### Spectroscopic investigations of recombinant RmPAE and triple mutant

#### Absorption and Fluorescence Spectroscopy

The absorption spectra for recombinant proteins of RmPAE and TM were recorded in the dark and after blue light illumination at 20 °C from 300 nm to 650 nm using Cary 3500 Compact UV-Vis Spectrophotometer equipped with peltier. Firstly, the absorption spectra of the dark-adapted recombinant proteins of RmPAE and TM were recorded. Further, the recombinant proteins of RmPAE and TM were irradiated with blue light for 30 seconds and the absorption spectra were again recorded after intermittent time intervals. Steady-state fluorescence emission spectra were also recorded for the same samples under similar conditions using Cary Eclipse Fluorescence Spectrometer. Emission spectra were recorded by exciting the protein samples at 390 nm and recording the emission from 400 nm to 700 nm.

#### Circular Dichroism Spectroscopy

The purified proteins were subjected to circular dichroism (CD) spectroscopy using a Jasco J-815 spectropolarimeter (JASCO, Japan). The spectra were obtained in the far-UV (200-250 nm) region at a concentration of 10 µM both in the dark and following blue light irradiation. The protein solution was in 10 mM sodium phosphate buffer containing 10 mM NaCl pH 7.4 in 1 mm path length quartz cuvette at temperature 20 °C. The instrument was equipped with a peltier and with N_2_ gas flowing. All the data were captured with the following settings: continuous mode, 50 nm/min scan speed, data pitch of 0.5 nm, sensitivity of 100 mdeg, response time of 8 s with 5 accumulations. In order to quantify the CD spectrum of the light-adapted condition, blue light illumination from an LED at 20 °C (luxeon ll Emitter LXHLPB09, 1 W, 460 nm; Conrad, Bavaria, Germany) during the CD measuring process. The sample was exposed to blue light for thirty seconds. All data were plotted after deducting the background signal from spectra by buffer solution’s spectrum as a reference.

#### Time resolved fluorescence spectroscopy

Time resolved fluorescence spectroscopy (TRFS) was performed to monitor the fluorescence decay of purified RmPAE and TM by using FLS1000 fluorescence spectrometer by Edinburgh Instruments. Pulsed laser diode and pulse width of 467 nm wavelength and 50 ps, respectively, was used for this study by employing the time correlated single photon counting (TCSPC). Fluorescence emission for RmPAE and TM was collected at 500 nm. Fluorescence decay curves and the Instrument Response Function (IRF) were plotted for both RmPAE and TM. The iterative re-convolution of IRF was done for fitting the graphs by using FAST software package by Edinburgh Instruments.

### Tertiary structure prediction and Molecular dynamic simulation RmPAE and TM

Tertiary structure for BLUF-EndoIII was predicted separately for BLUF and Endonuclease domains using alpha-fold 3 protein structure prediction server (Abramson et al. 2024; https://alphafoldserver.com/welcome). The predicted structures were validated using SAVES server (https://saves.mbi.ucla.edu/) (Pontius et al. 1996). Predicted structure were visualized using Discovery Studio v24. The molecular dynamics (MD) simulation was performed using GROMACS V2022 software (Abraham et al. 2015; Kutzner et al. 2007), to understand the effect of mutation on FMN binding ability with the BLUF domain from RmPAE and TM (BLUF-EndoIII Y5F, H27N, W87A). The three-dimensional (3D) structures of RmPAE and TM were predicted using alpha-fold. The simulations were performed at 300K temperature for 100 ns with water as the solvent. AMBER (Assisted Model Building and Energy Refinement) 99SB force field was implemented for the simulations and 1.2 nm protein to box distance was used (Case et al. 2023; Maier et al. 2015; Lindorff-Larsen et al. 2010; Hornak et al. 2006; Cornell et al. 1995). The systems were solvated using the TIP3P water molecules and the counter ions were used to neutralized the system (William et al. 1983). Both van der Waals cut-off and electrostatic value were set at1.2 nm, using the Particle Mesh Ewald algorithm (Essmann et al. 1995). The two-step ensemble techniques, NVT and NPT, were used to equilibrate the systems for 100 nm. For the NVT canonical ensemble, the Berendsen thermostat without pressure coupling was run at 300K (Bussi et al. 2007; Berendsen et al. 1984). The steepest descent method was used to minimize the entire system until the energy value was less than 10 kJ/Mol. MD simulations for all systems were performed for 100 ns using the leapfrog integrator to track the evolution of run paths. VMD (Visual molecular dynamics) and PyMOL software were used to create the graphical representations. Data analysis was done through Gromacs and generate by gnu plot.

### Endonuclease assay of purified RmPAE and triple mutant

The endonuclease activity of BLUF-EndoIII was carried out with different substrates under dark and blue light conditions. Purified recombinant proteins (11 µg/µl) were incubated at 37 °C for 30 minutes with substrate plasmid DNA (pGEX; 300 ng/µl) in absence (dark) and presence of blue light (BL). The pGEX vector alone was used as a control. Reactions were stopped with DNA loading dye containing Tris-Cl (pH = 8) and were separated on agarose gel. All the samples were separated on 1% electrophoretic gel.

### Protein-DNA interaction analysis

#### In-silico prediction of DNA-Protein Interactions

The protein-DNA interaction analysis was performed using HDOCK server (http://hdock.phys.hust.edu.cn/; Yan et al. 2020). The 500 bp segments of pGEX vector sequence (4932 bps) were used to prepare the DNA molecules (in PDB format). Further, docking analysis of TM (receptor molecule) was performed separately with each of the DNA fragments (ligand molecule). The output of each docking analysis presents top 10 docked complex; out of which, first complex (Rank 1) with maximum negative docking score and confidence score of ∼0.7 was considered the best docking complex. A more negative docking score and confidence score of ∼ 0.7 suggested that the two input molecules would be very likely to bind. The interaction in the predicted complex were visualized using Discovery Studio v24. The interaction between TM and the *Roseophage* SI01DNA fragments were also performed. We have taken starting 3kbp sequence of *Roseophage* SI01DNA (∼40.5 kbp) and used 500 bp segments to prepare the DNA molecules (in PDB format) and employed each segment for docking analysis against TM (receptor molecule). The top model complex with maximum negative docking score and confidence score of ∼0.7 was visualized using Discovery Studio v24 to predicting interacting partners.

#### DNA-Protein Interactions Analysis by Intrinsic Fluorescence estimation

The interaction between the protein and predicted oligonucleotide was estimated by monitoring the change in the fluorescence intensity of free and oligonucleotide-bound protein. The predicted oligonucleotide fragments were commercially synthesised and employed for the DNA-protein interaction analysis. The dilutions of recombinant TM and oligonucleotides were prepared in the interaction buffer [HEPES buffer (20 mM; pH 7.4), MgCl_2_ (20 mM), potassium acetate (100 mM), DTT (2 mM), 0.2% Tween-20, 5% glycerol, BSA (0.1 mg ml^−1^)] and the concentration were estimated using standard protocol. Further, the fluorescence emission spectrum (scan speed will be set at 60 nm min^−1^) of the TM alone was estimated by exciting the protein samples at 390 nm and recording the emission from 400 nm to 700 nm. After monitoring the emission spectra for protein alone, add small aliquot of oligonucleotide solution, mix and immediately check the fluorescence emission at the emission maximum of the protein. Gradually increase the volume of the oligonucleotide and monitor the change in fluorescence intensity at each point. Continue until no change in change in fluorescence intensity was observed for several points. The oligonucleotide concentration was also calculated at each point using following equation:

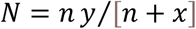

Where, n is the total volume of oligonucleotide added, x is the initial volume of TM added, and y is the molar concentration of oligonucleotide stock solution.

The DNA-protein interaction dynamics was studied by using stern-volmer analysis which shows the reduction in the fluorescence intensity as the function of quencher (oligonucleotide) concentration (Q). The stern-volmer equation is described as:

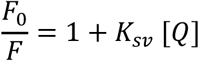

Where, F_0_ and F is the fluorescence intensity of free and oligonucleotide-bound TM, respectively; K_sv_ is stern-volmer quenching constant and Q is the concentration of quencher/oligonucleotide.

Further, the biomolecular quenching rate constant (K_q_) for DNA-protein interaction was estimated by using following equation:

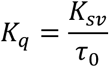

Where τ_0_ is the excited state fluorescence lifetime of protein (TM) in the absence of quencher (oligonucleotide).

## Author Contributions

Rahamtullah and Jitender had performed the biochemical characterization of the recombinant proteins and done the structure function analysis. Rahamtullah, Jitender and MSK had performed the bioinformatics and MD simulation related analysis. MSK and Jitender had done the DNA-protein interaction analysis. Sagar Kumar had performed the SDM analysis. S.K. had conceived the project. S.K. and MSK wrote the manuscript.

## Acknowledgments

Rahmatullah and M.S.K. were supported from 52/310/2022-BIO-BMS-JNU. Jitender was supported by CSIR-JRF. S.K. was supported from STARS/APR2019/BS/563/FS and SPARC/2019-2020/P2127/SL. We also acknowledge CRF, IIT-Delhi and CIF, JNU for providing research facility.

## Data availability

All data supporting the findings of this study are available in the article and its supplementary information section.

## Extended Data Figures and Figure Legends

**Table S1.** Time constant calculated from the dark state recovery of the BLUF Endonuclease III wild type and mutant proteins. (BE-BLUF Endonuclease III). ND-not determined.

| Sample | Absorption maximum (nm) | Time constant, $\tau$ (sec) |
| --- | --- | --- |
| RmPAE | 440 | 462 |
| Triple mutant (TM) | 440 | 687 |

**Table S2.** Life time and fractional intensity contributions of above proteins. χ^2^ represents goodness of fit), B1, B2 and B3 represents the fractional intensity contribution of each component, τ_i_ represents the life-time for each component, <τ> represents the average life-time. (BE-BLUF Endonuclease III

| Sample | $\chi^2$ | B1 | $\tau_1$ (ns) | B2 | $\tau_2$ (ns) | B3 | $\tau_3$ (ns) | $\langle\tau\rangle$ |
| --- | --- | --- | --- | --- | --- | --- | --- | --- |
| RmPAE | 1.14 | 0.03 | 0.50 | 0.01 | 4.59 | 0.96 | 0.067 | 0.13 |
| Triple mutant (TM) | 1.07 | 0.50 | 1.96 | 0.31 | 3.08 | 0.182554 | 0.33 | 2.02 |

**Table S3.**
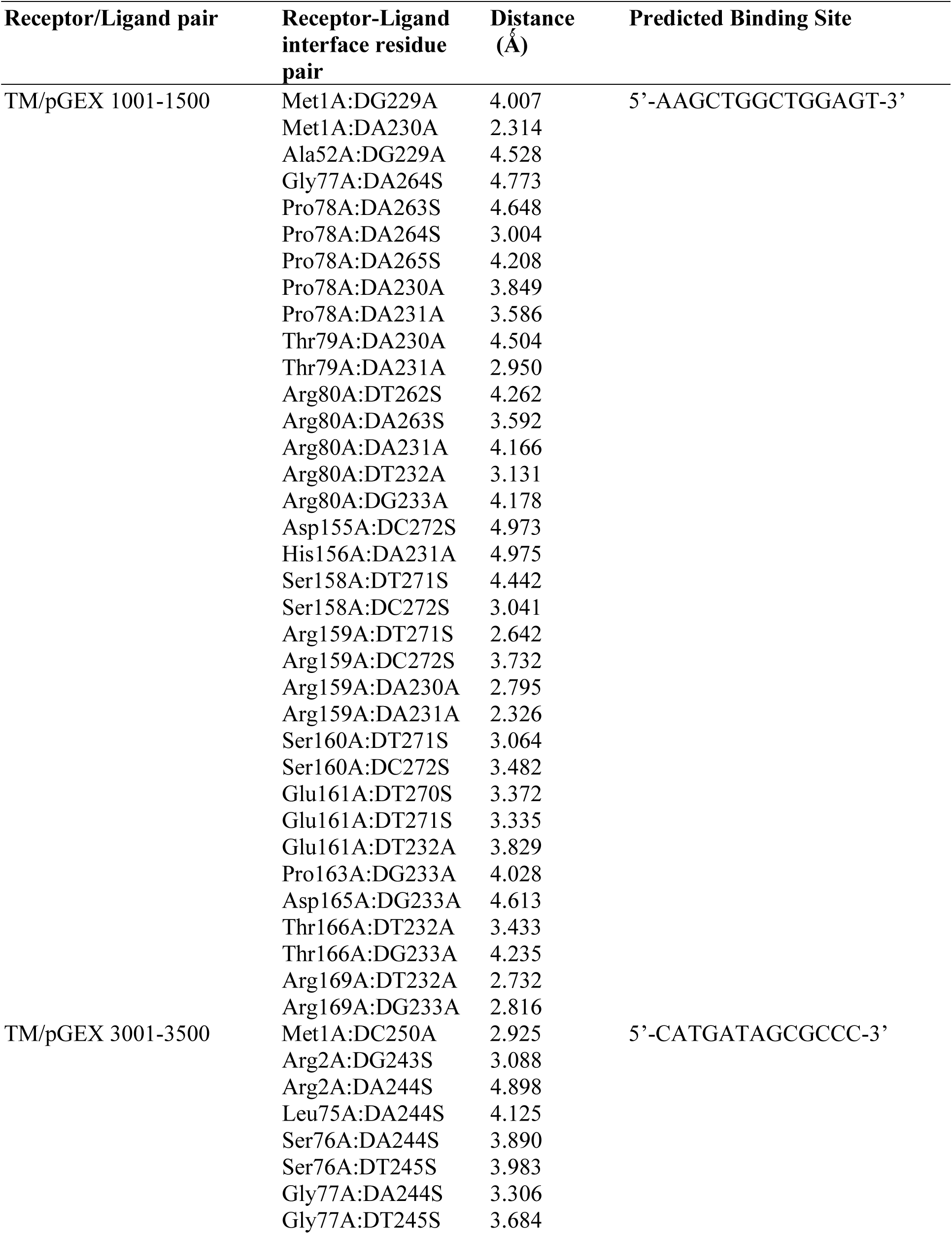
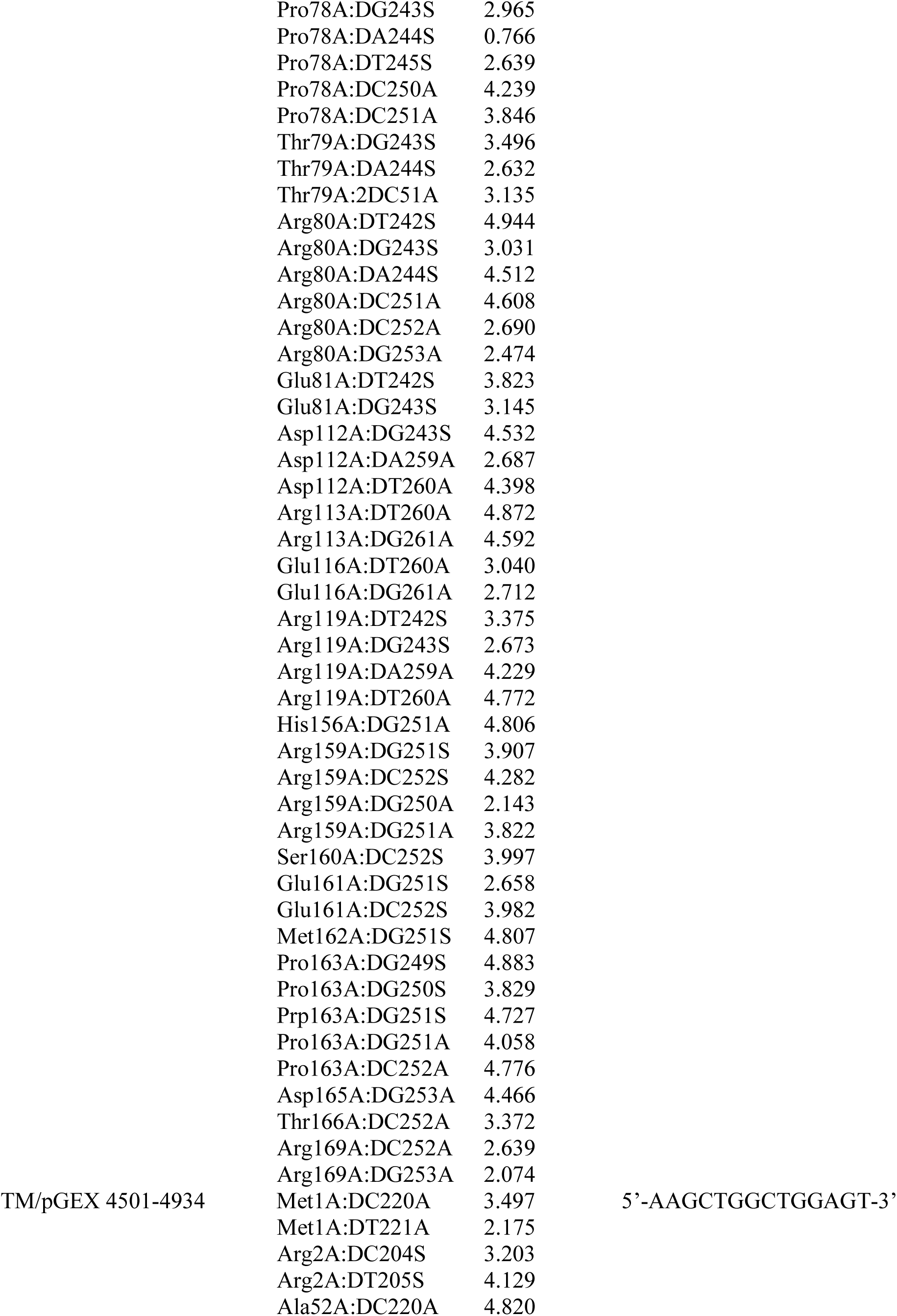
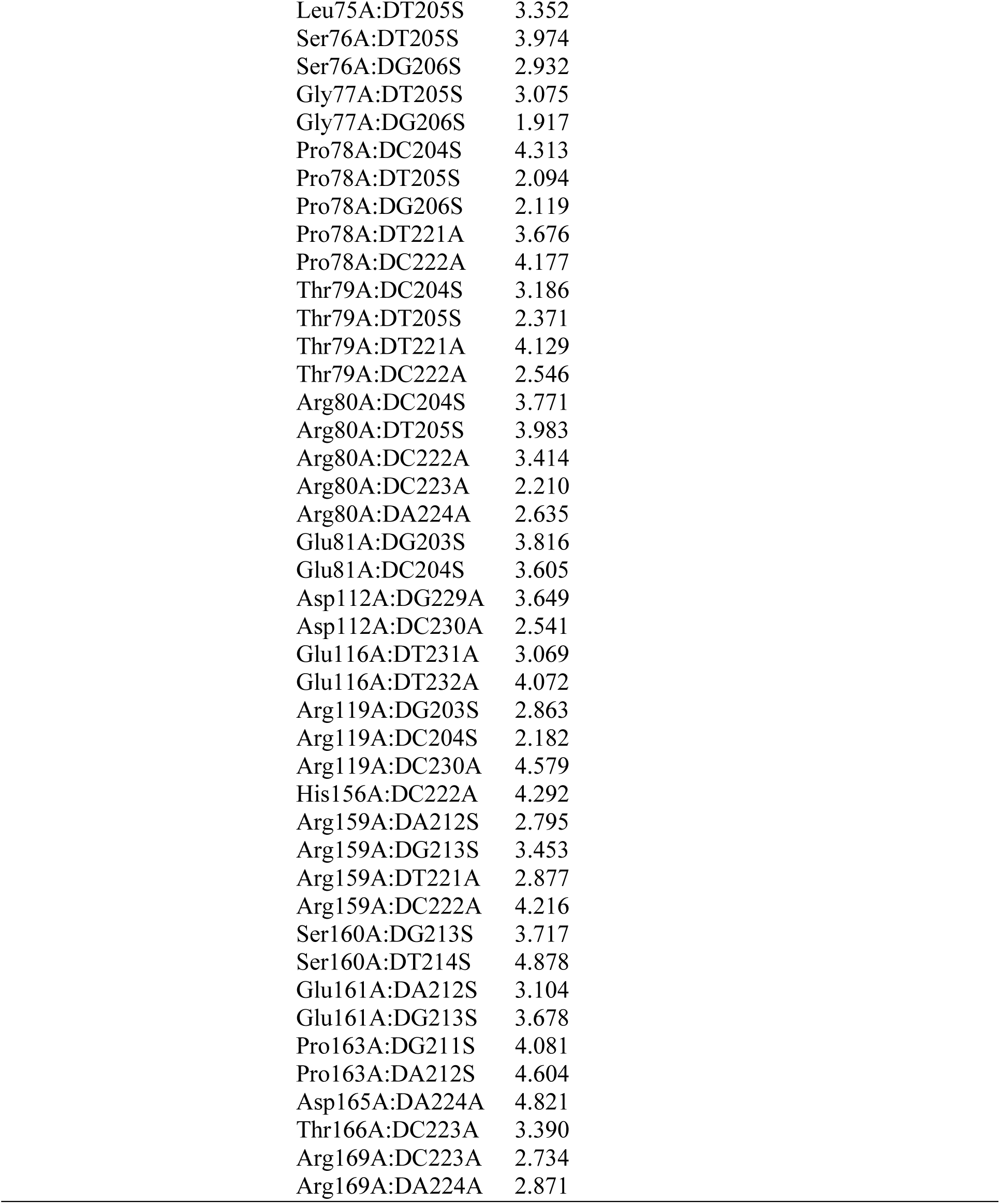
Interaction output of docking analysis of TM and fragments of pGEX vector (1001-1500 bps, 3001-3500 bps, and 4501-4934 bp) as receptor and ligand molecules, respectively.

| Receptor/Ligand pair | Receptor-Ligand interface residue pair | Distance (Å) | Predicted Binding Site |
| --- | --- | --- | --- |
| TM/pGEX 1001-1500 | Met1A:DG229A | 4.007 | 5'-AAGCTGGCTGGAGT-3' |
|  | Met1A:DA230A | 2.314 |  |
|  | Ala52A:DG229A | 4.528 |  |
|  | Gly77A:DA264S | 4.773 |  |
|  | Pro78A:DA263S | 4.648 |  |
|  | Pro78A:DA264S | 3.004 |  |
|  | Pro78A:DA265S | 4.208 |  |
|  | Pro78A:DA230A | 3.849 |  |
|  | Pro78A:DA231A | 3.586 |  |
|  | Thr79A:DA230A | 4.504 |  |
|  | Thr79A:DA231A | 2.950 |  |
|  | Arg80A:DT262S | 4.262 |  |
|  | Arg80A:DA263S | 3.592 |  |
|  | Arg80A:DA231A | 4.166 |  |
|  | Arg80A:DT232A | 3.131 |  |
|  | Arg80A:DG233A | 4.178 |  |
|  | Asp155A:DC272S | 4.973 |  |
|  | His156A:DA231A | 4.975 |  |
|  | Ser158A:DT271S | 4.442 |  |
|  | Ser158A:DC272S | 3.041 |  |
|  | Arg159A:DT271S | 2.642 |  |
|  | Arg159A:DC272S | 3.732 |  |
|  | Arg159A:DA230A | 2.795 |  |
|  | Arg159A:DA231A | 2.326 |  |
|  | Ser160A:DT271S | 3.064 |  |
|  | Ser160A:DC272S | 3.482 |  |
|  | Glu161A:DT270S | 3.372 |  |
|  | Glu161A:DT271S | 3.335 |  |
|  | Glu161A:DT232A | 3.829 |  |
|  | Pro163A:DG233A | 4.028 |  |
|  | Asp165A:DG233A | 4.613 |  |
|  | Thr166A:DT232A | 3.433 |  |
|  | Thr166A:DG233A | 4.235 |  |
|  | Arg169A:DT232A | 2.732 |  |
|  | Arg169A:DG233A | 2.816 |  |
| TM/pGEX 3001-3500 | Met1A:DC250A | 2.925 | 5'-CATGATAGCGCCC-3' |
|  | Arg2A:DG243S | 3.088 |  |
|  | Arg2A:DA244S | 4.898 |  |
|  | Leu75A:DA244S | 4.125 |  |
|  | Ser76A:DA244S | 3.890 |  |
|  | Ser76A:DT245S | 3.983 |  |
|  | Gly77A:DA244S | 3.306 |  |
|  | Gly77A:DT245S | 3.684 |  |

|  |  |  |  |  |
| --- | --- | --- | --- | --- |
| 1044 |  | Pro78A:DG243S | 2.965 |  |
| 1045 |  | Pro78A:DA244S | 0.766 |  |
| 1046 |  | Pro78A:DT245S | 2.639 |  |
| 1047 |  | Pro78A:DC250A | 4.239 |  |
| 1048 |  | Pro78A:DC251A | 3.846 |  |
| 1049 |  | Thr79A:DG243S | 3.496 |  |
| 1050 |  | Thr79A:DA244S | 2.632 |  |
| 1051 |  | Thr79A:2DC51A | 3.135 |  |
| 1052 |  | Arg80A:DT242S | 4.944 |  |
| 1053 |  | Arg80A:DG243S | 3.031 |  |
| 1054 |  | Arg80A:DA244S | 4.512 |  |
| 1055 |  | Arg80A:DC251A | 4.608 |  |
| 1056 |  | Arg80A:DC252A | 2.690 |  |
| 1057 |  | Arg80A:DG253A | 2.474 |  |
| 1058 |  | Glu81A:DT242S | 3.823 |  |
| 1059 |  | Glu81A:DG243S | 3.145 |  |
| 1060 |  | Asp112A:DG243S | 4.532 |  |
| 1061 |  | Asp112A:DA259A | 2.687 |  |
| 1062 |  | Asp112A:DT260A | 4.398 |  |
| 1063 |  | Arg113A:DT260A | 4.872 |  |
| 1064 |  | Arg113A:DG261A | 4.592 |  |
| 1065 |  | Glu116A:DT260A | 3.040 |  |
| 1066 |  | Glu116A:DG261A | 2.712 |  |
| 1067 |  | Arg119A:DT242S | 3.375 |  |
| 1068 |  | Arg119A:DG243S | 2.673 |  |
| 1069 |  | Arg119A:DA259A | 4.229 |  |
| 1070 |  | Arg119A:DT260A | 4.772 |  |
| 1071 |  | His156A:DG251A | 4.806 |  |
| 1072 |  | Arg159A:DG251S | 3.907 |  |
| 1073 |  | Arg159A:DC252S | 4.282 |  |
| 1074 |  | Arg159A:DG250A | 2.143 |  |
| 1075 |  | Arg159A:DG251A | 3.822 |  |
| 1076 |  | Ser160A:DC252S | 3.997 |  |
| 1077 |  | Glu161A:DG251S | 2.658 |  |
| 1078 |  | Glu161A:DC252S | 3.982 |  |
| 1079 |  | Met162A:DG251S | 4.807 |  |
| 1080 |  | Pro163A:DG249S | 4.883 |  |
| 1081 |  | Pro163A:DG250S | 3.829 |  |
| 1082 |  | Prp163A:DG251S | 4.727 |  |
| 1083 |  | Pro163A:DG251A | 4.058 |  |
| 1084 |  | Pro163A:DC252A | 4.776 |  |
| 1085 |  | Asp165A:DG253A | 4.466 |  |
| 1086 |  | Thr166A:DC252A | 3.372 |  |
| 1087 |  | Arg169A:DC252A | 2.639 |  |
| 1088 |  | Arg169A:DG253A | 2.074 |  |
| 1089 | TM/pGEX 4501-4934 | Met1A:DC220A | 3.497 | 5'-AAGCTGGCTGGAGT-3' |
| 1090 |  | Met1A:DT221A | 2.175 |  |
| 1091 |  | Arg2A:DC204S | 3.203 |  |
| 1092 |  | Arg2A:DT205S | 4.129 |  |
| 1093 |  | Ala52A:DC220A | 4.820 |  |

|  |  |  |
| --- | --- | --- |
| 1094 | Leu75A:DT205S | 3.352 |
| 1095 | Ser76A:DT205S | 3.974 |
| 1096 | Ser76A:DG206S | 2.932 |
| 1097 | Gly77A:DT205S | 3.075 |
| 1098 | Gly77A:DG206S | 1.917 |
| 1099 | Pro78A:DC204S | 4.313 |
| 1100 | Pro78A:DT205S | 2.094 |
| 1101 | Pro78A:DG206S | 2.119 |
| 1102 | Pro78A:DT221A | 3.676 |
| 1103 | Pro78A:DC222A | 4.177 |
| 1104 | Thr79A:DC204S | 3.186 |
| 1105 | Thr79A:DT205S | 2.371 |
| 1106 | Thr79A:DT221A | 4.129 |
| 1107 | Thr79A:DC222A | 2.546 |
| 1108 | Arg80A:DC204S | 3.771 |
| 1109 | Arg80A:DT205S | 3.983 |
| 1110 | Arg80A:DC222A | 3.414 |
| 1111 | Arg80A:DC223A | 2.210 |
| 1112 | Arg80A:DA224A | 2.635 |
| 1113 | Glu81A:DG203S | 3.816 |
| 1114 | Glu81A:DC204S | 3.605 |
| 1115 | Asp112A:DG229A | 3.649 |
| 1116 | Asp112A:DC230A | 2.541 |
| 1117 | Glu116A:DT231A | 3.069 |
| 1118 | Glu116A:DT232A | 4.072 |
| 1119 | Arg119A:DG203S | 2.863 |
| 1120 | Arg119A:DC204S | 2.182 |
| 1121 | Arg119A:DC230A | 4.579 |
| 1122 | His156A:DC222A | 4.292 |
| 1123 | Arg159A:DA212S | 2.795 |
| 1124 | Arg159A:DG213S | 3.453 |
| 1125 | Arg159A:DT221A | 2.877 |
| 1126 | Arg159A:DC222A | 4.216 |
| 1127 | Ser160A:DG213S | 3.717 |
| 1128 | Ser160A:DT214S | 4.878 |
| 1129 | Glu161A:DA212S | 3.104 |
| 1130 | Glu161A:DG213S | 3.678 |
| 1131 | Pro163A:DG211S | 4.081 |
| 1132 | Pro163A:DA212S | 4.604 |
| 1133 | Asp165A:DA224A | 4.821 |
| 1134 | Thr166A:DC223A | 3.390 |
| 1135 | Arg169A:DC223A | 2.734 |
| 1136 | Arg169A:DA224A | 2.871 |
| 1137 |  |  |
| 1138 |  |  |
| 1139 |  |  |
| 1140 |  |  |
| 1141 |  |  |
| 1142 |  |  |
| 1143 |  |  |

**Table S4.**
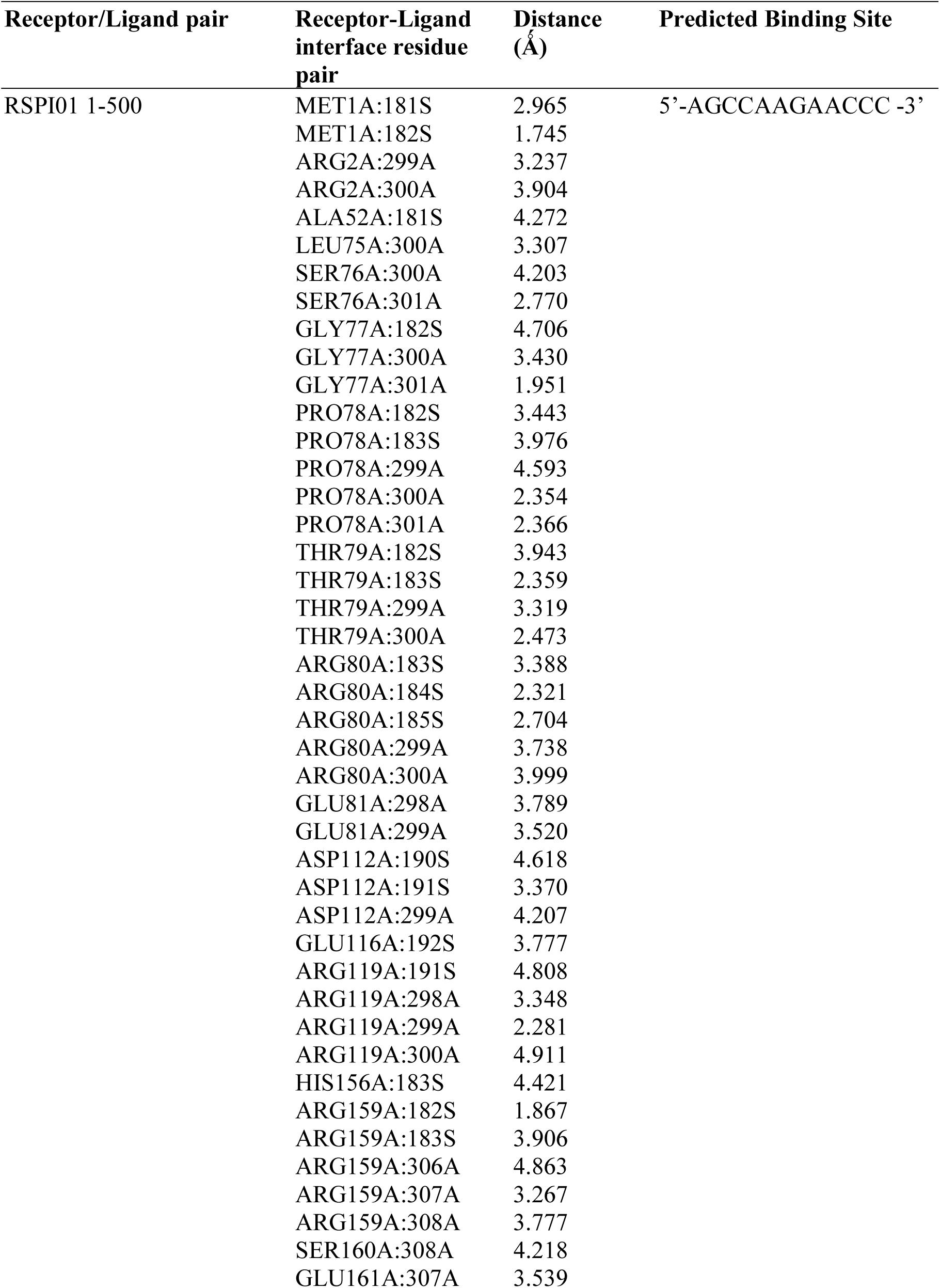
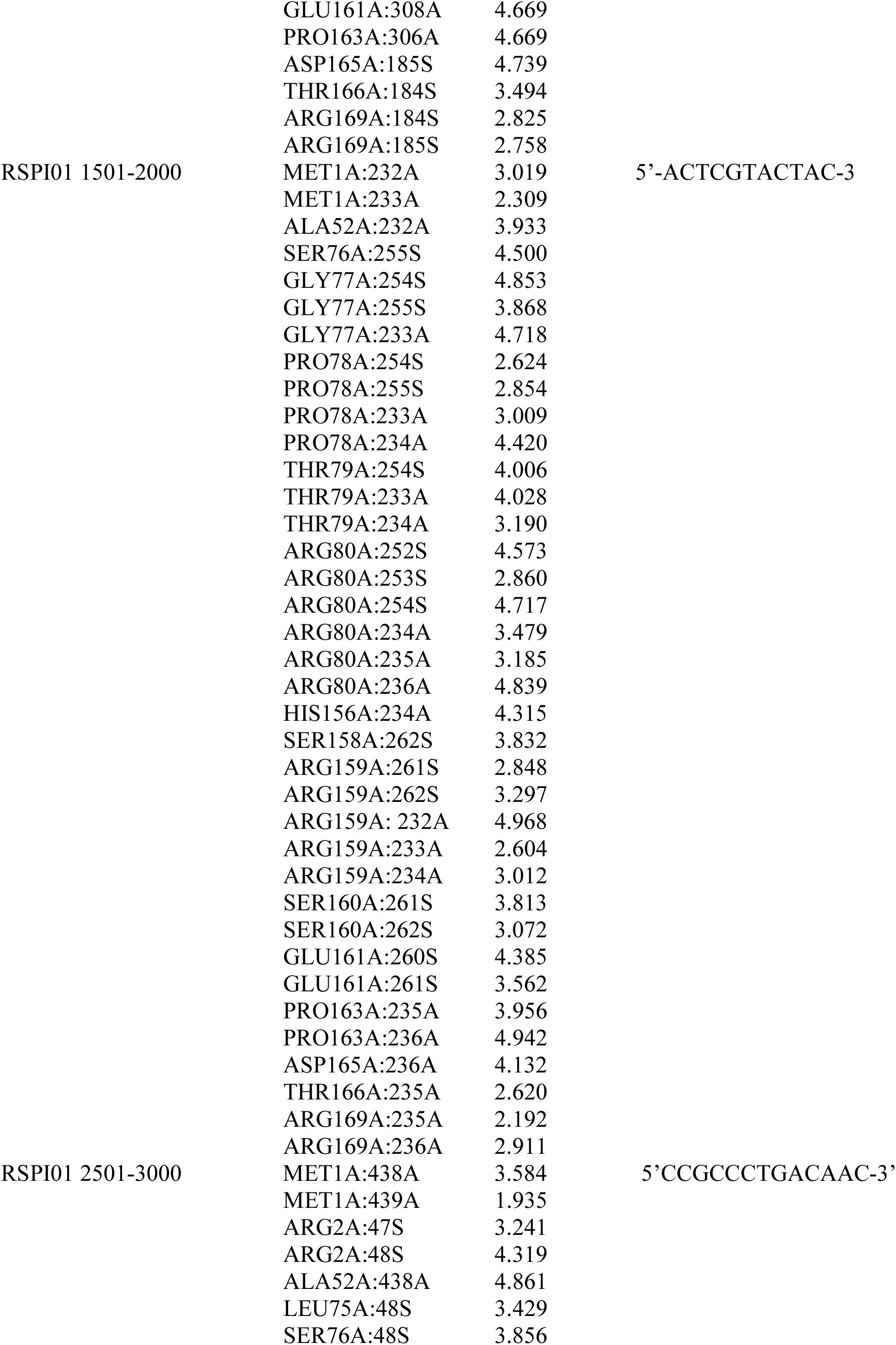
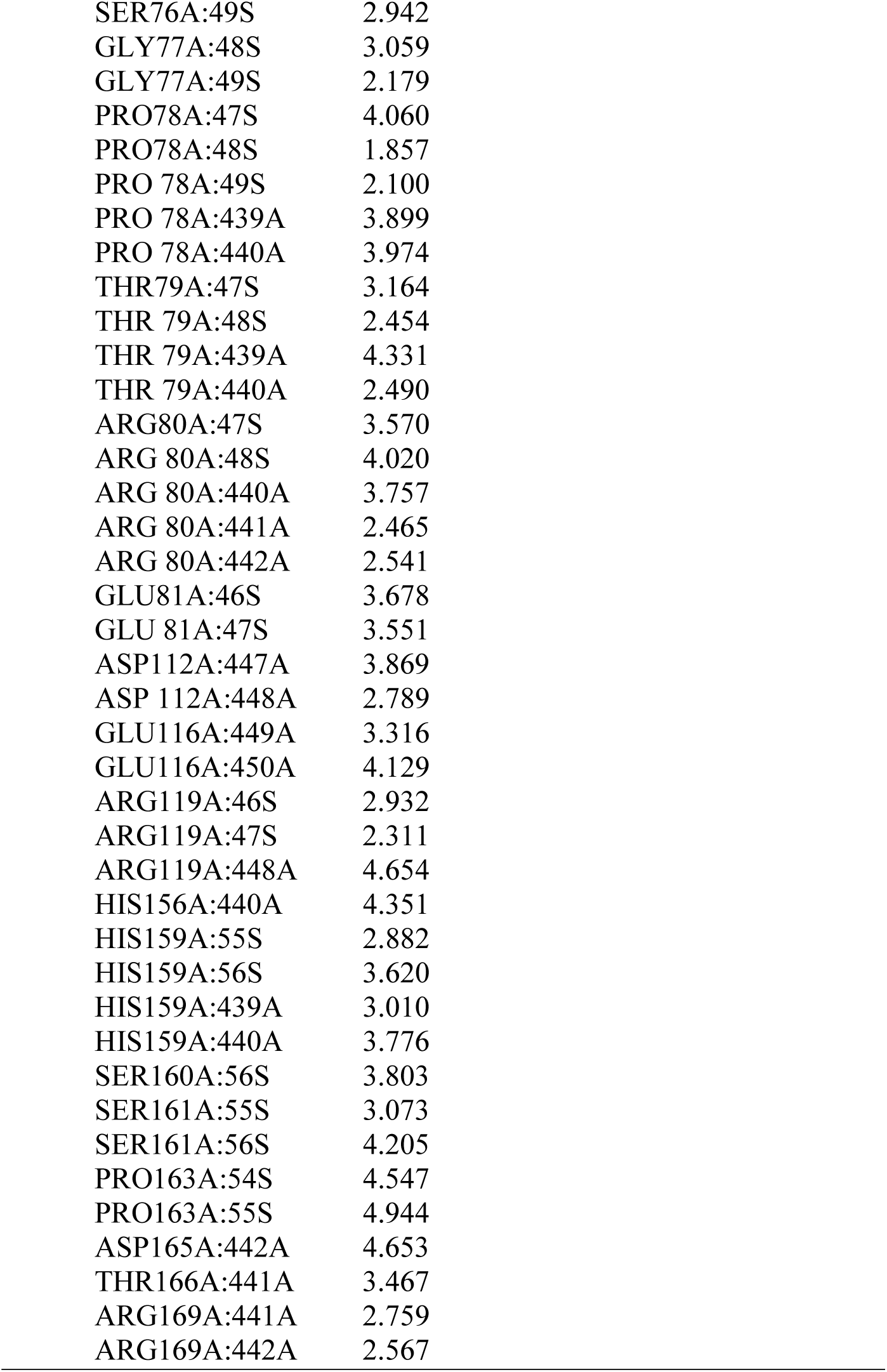
Interaction output of docking analysis of TM and fragments of *Roseophage* SI01 (1-500 bps, 1501-2000 bps, and 2501-3000 bp) as receptor and ligand molecules, respectively.

**Fig. S1.**
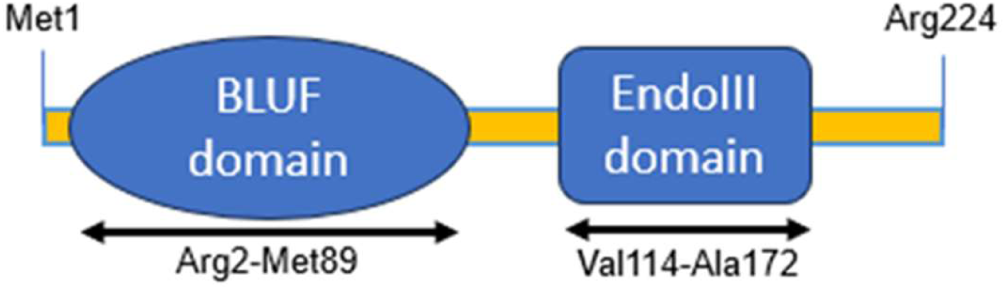
Diagrammatic representation of *R. mesophilum’s* BLUF-EndoIII domain organization. The BLUF domain ranges from 2-89 amino acids and the endonuclease III domain is from 114–172 amino acids.

**Fig. S2.**
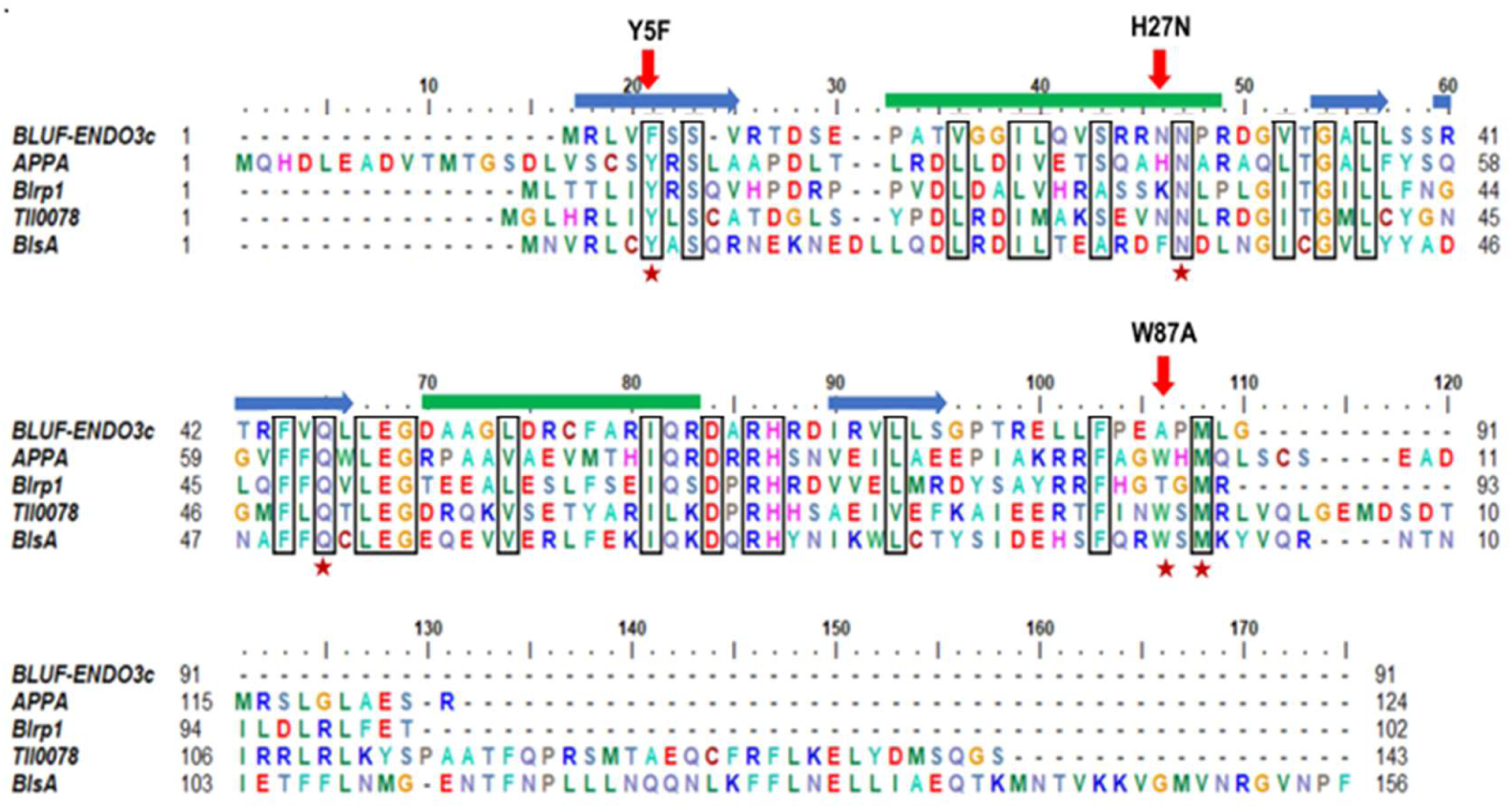
Multiple sequence alignment of BLUF domain from the RmPAE, BLUF-EndoIII from *R. mesophilum*, AppA from *Rhodobacter sphaeroides,* Blrp1 from *Klebsiella pneumoniae*, TII0078 from *Thermosynechococcus vestitus*, and BlsA from *Acinetobacter baumannii*. Red arrow indicates positions of mutated amino acids (Y5F, H27N, W87A). Asterisks (*) represents amino acids crucial for BLUF photocycle. Dark blue horizontal arrows indicate α-helices and green horizontal thick lines represent β-sheets.

**Fig. S3.**
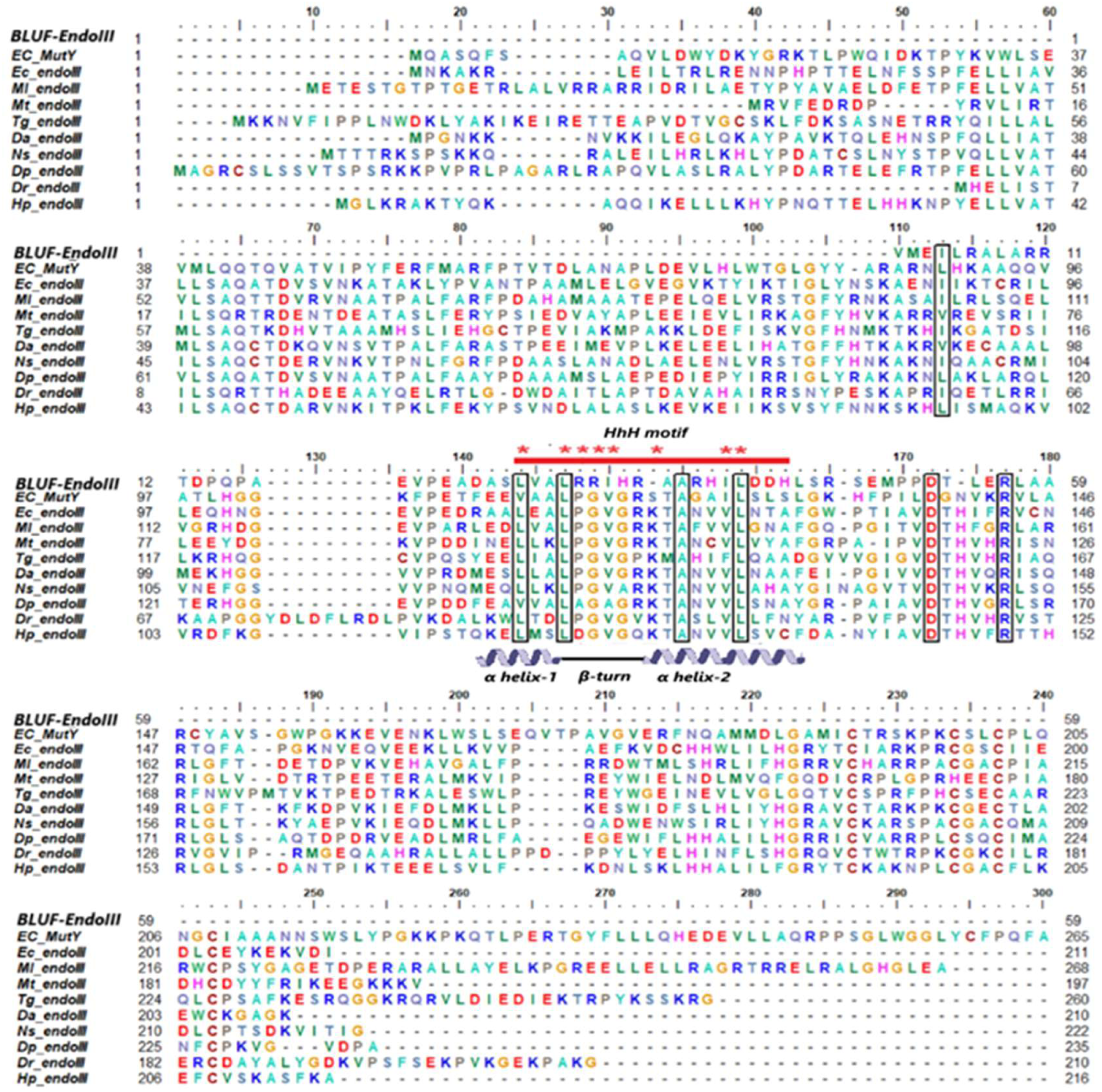
Multiple sequence alignment of BLUF-EndoIII. Sequence alignment of Endonuclease domain of RmPAE and BLUF-EndoIII from *R.* mesophilum against well characterized MutY from *E. coli*, endonuclease III domains from *E. coli, Micrococcus luteus*, and *Methanothermobacter thermautotrophicus.* Alignment shows in the presence of the characteristic HtH motif (indicated with red solid line). Aspartate residue (D52), an active site residue is highly conserved in all members of endonuclease III family. Amino acids important for the endonuclease activity are marked with asterisk. (BLUF-EndoIII-*Rubellimicrobium mesophilum* endonuclease III, Ec EndoIII-*Escherichia coli* endonuclease III, Ec MutY-*Escherichia coli* MutY, Ml EndoIII-*Micrococcus luteus* endonuclease III, Mt EndoIII-*Methanothermobacter thermautotrophicus* endonuclease III and RmPAE Endo-endonuclease domain from synthetic BLUF-EndoIII).

**Fig. S4.**
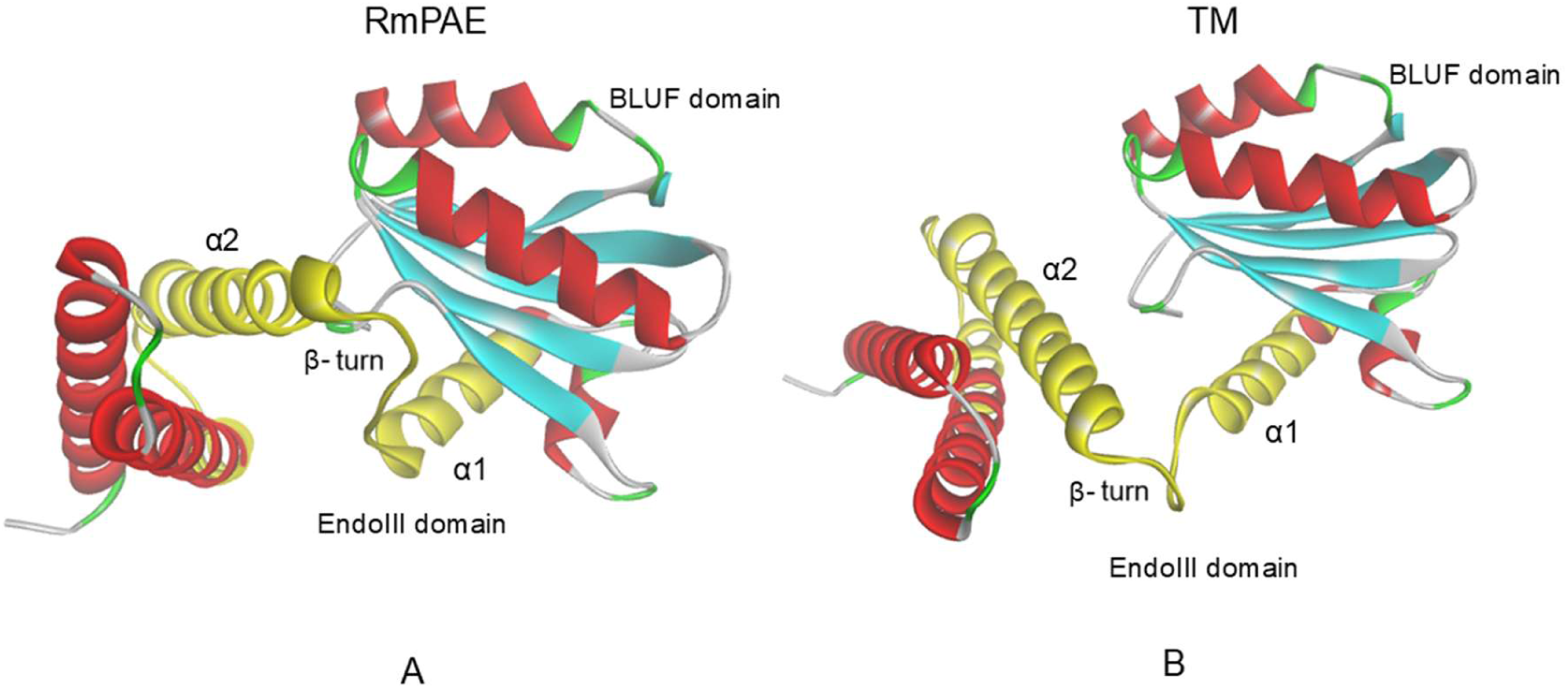
Tertiary structures of RmPAE (A) and TM (B) predicted using alpha-fold3 protein structure prediction server and visualized using discovery studio v24. In both RmPAE and Tm, BLUF domain consists of five-stranded β-sheet flanked by two α-helices. The EndoIII domain is presented in yellow colour and consist of a helix-hairpin-helix (HhH) motif composed α-helix 1 (α1) and α-helix 2 (α2)which is separated by a β-turn.

**Fig. S5.**
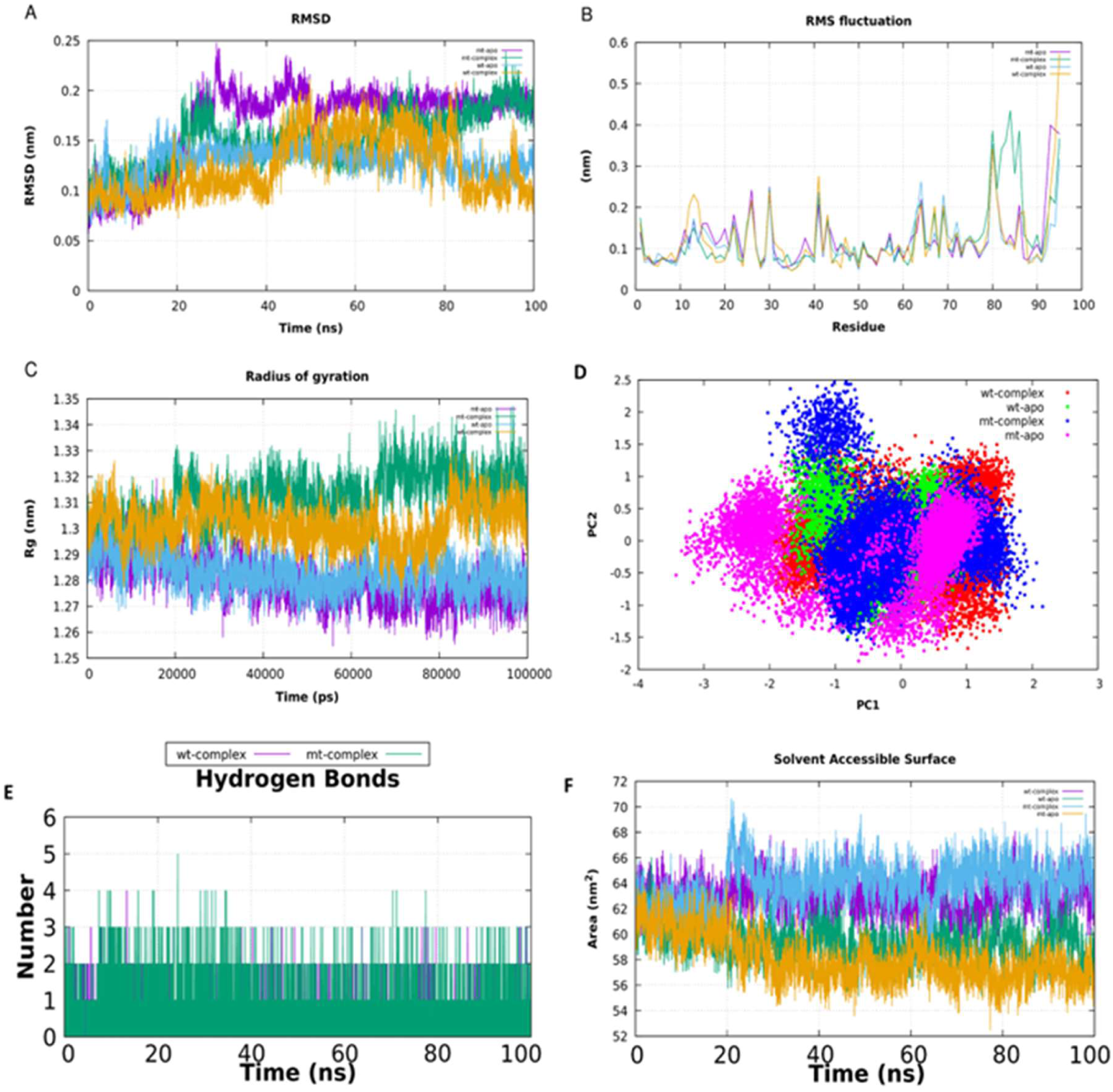
The impact of the mutation on FMN binding is investigated using simulation studies. (A) RMSD (Root Mean Square Deviation), (B) RMS Fluctuation (C) Radius of gyration (Rg) (D) Principal component analysis; different form is indicated in different colours, wild type apo, wild type-complex, mutant apo and mutant complex are indicated as green, red, pink and blue respectively. (E) Hydrogen bond, Time evolution of hydrogen bonds formed between BLUF endonuclease proteins and FMN (F) SASA (Solvent Accessibility Surface Area). In the figure A,B,C,E and F different form of protein is indicated as different colour, wild type apo, wild type-complex, mutant apo and mutant complex are indicated as blue, yellow, magenta and green respectively.

